# ULK1-linked mitophagy promotes cardiac hypoxia tolerance in the blind mole-rat

**DOI:** 10.64898/2026.03.13.711419

**Authors:** Damla Yalcin, Sudenaz Fatma Oner, Erdogan Oguzhan Akyildiz, Faruk Colak, Cigdem Kayra Ongun, Hsin-Yu Fang, Ismail Selim Yıldırım, Preeti Bais, Thomas J. Park, Emel Timucin, Sukru Anil Dogan, Erkan Tuncay, Perinur Bozaykut

## Abstract

Blind mole-rats (BMRs) thrive in chronically hypoxic subterranean environments, displaying exceptional cardiac resilience to conditions that rapidly induce failure in other mammals. Here, we integrate *in vivo* physiology, multi-omics profiling, mitochondrial analyses, and genome editing to uncover an evolved cardioprotective program in BMRs. Under acute 0% O₂ exposure, BMRs exhibit markedly prolonged survival compared to mouse. At the molecular level, BMR hearts undergo coordinated metabolic remodeling, restrained inflammatory signaling, and enhanced genome maintenance. Functionally, BMR cardiac mitochondria suppress high-flux oxidative phosphorylation and reverse electron transport–associated ROS following hypoxia, indicating intrinsic adaptation to oxygen collapse. Hypoxia selectively activates AMPK–mTOR–ULK1 dependent mitophagy, and pharmacological manipulation demonstrates that mitophagy is required for BMR cardiomyocyte survival during hypoxia-reoxygenation stress. Finally, we identify a BMR-specific insertion in ULK1 and demonstrate that introduction of this sequence into rat cardiomyocytes enhances hypoxia tolerance in a mitophagy-dependent manner. These findings reveal an evolutionarily tuned mitochondrial quality-control strategy that enables extreme cardiac resilience to hypoxia.

## Introduction

Most mammals have limited capacity to tolerate environmental extremes, particularly oxygen deprivation, which rapidly disrupts mitochondrial function, increases reactive oxygen species (ROS) production, and induces tissue damage [1,2]. Even brief interruptions in oxygen supply destabilize oxidative phosphorylation (OXPHOS) in the heart, triggering metabolic collapse and tissue injury [3,4]. In contrast, a select group of naturally hypoxia-adapted mammals display a remarkable capacity to withstand extreme oxygen limitation and provide powerful comparative models for uncovering endogenous mechanisms of tissue protection [5,6].

The blind mole-rat (BMR; *Spalax/Nannospalax sp*) represents one of the most extreme example of mammalian adaptation to chronic hypoxia. Living in subterranean burrow systems with chronically low and fluctuating oxygen levels, BMRs exhibit striking phenotypic traits, including pronounced hypoxia tolerance together with cancer resistance, delayed aging, and exceptional longevity that imply deep evolutionary remodeling of physiological stress-response networks [7–12]. Such adaptations suggest that BMRs sustain protective programs that are absent in hypoxia-sensitive species, particularly in metabolically demanding organs such as the heart.

Mitochondria are central determinants of cardiomyocyte survival during oxygen stress [13]. While OXPHOS is essential for ATP production, it also constitutes a major source of ROS, especially under conditions that promote reverse electron transport (RET) at complex I [14]. Excess ROS exacerbates mitochondrial dysfunction and activates cell-death pathways [15–17]. Mitochondrial quality-control mechanisms, particularly mitophagy, counteract this process by selectively eliminating damaged organelles, stabilizing ATP supply and limiting ROS accumulation [18–21]. These pathways are subject to upstream regulation by nutrient sensing kinases, AMP-activated protein kinase (AMPK) and the mechanistic target of rapamycin (mTOR) [22–25]. Under metabolic stress, AMPK activation suppresses mTOR signalling and promotes autophagy through activation of UNC-51–like kinase 1 (ULK1), a serine/threonine kinase that initiates autophagosome formation [26]. Hypoxia can activate this regulatory circuit through the AMPK–mTOR–ULK1 axis, and pharmacological stimulation of this pathway has been shown to attenuate hypoxic injury [27]. Despite extensive characterization of these conserved energy-sensing and mitochondrial-quality control pathways in traditional model organisms, it remains unknown how they are tuned in naturally hypoxia-tolerant mammals and whether evolutionary pressures have modified these networks to confer enhanced cardiac resilience.

Here, we address this gap by leveraging the BMR as a comparative model to define an evolutionarily shaped cardioprotective program. By integrating *in vivo* hypoxia physiology with multi-omics profiling, mitochondrial functional analyses, and cell-intrinsic hypoxia–reoxygenation assays, we define a coordinated hypoxia response that suppresses oxygen-intensive mitochondrial states while selectively engaging mitophagy.

We further uncover a lineage-specific insertion within the ULK1 kinase domain and establish causality using CRISPR-mediated knock-in in rat cardiomyocytes, demonstrating that this sequence variant enhances survival following hypoxia–reoxygenation. Together, these results reveal how evolutionary adaptation can reconfigure conserved stress-response pathways to achieve extreme hypoxia tolerance in a long-lived mammalian heart.

## Results

### Blind mole-rats exhibit enhanced physiological resilience to severe hypoxia

To characterize species-specific mechanisms of hypoxia resilience, blind mole-rats (BMR; *Nannospalax* spp.) and BALB/c mice (*Mus musculus*) were subjected to extreme oxygen deprivation (0% O₂) using a controlled hypoxia paradigm adapted from prior work in naked mole-rats [28]. Animals were exposed to hypoxia for empirically determined durations (mouse, 300 s; BMR, 650 s) selected to encompass the terminal physiological failure window for each species, followed by collection of cardiac and lung tissues for downstream analyses (Fig. 1A).

**Fig 1.**
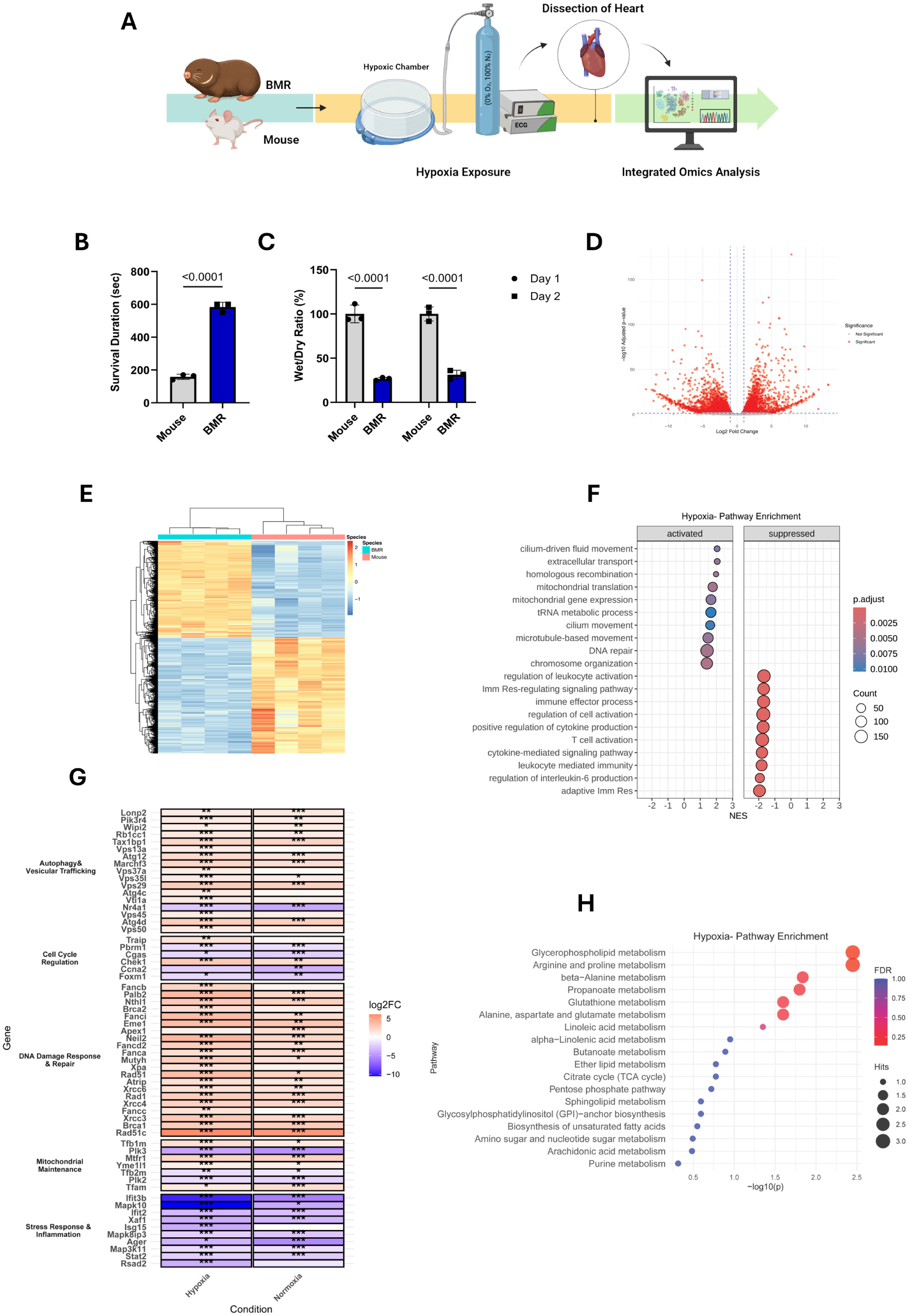
Extreme hypoxia tolerance in BMRs is coupled with cardiac multi-omic remodeling. **(A)** Schematic overview of the *in vivo* hypoxia paradigm and downstream analyses. Blind mole-rats (BMRs) and mouse were exposed to normoxia (∼21% O₂) or acute severe hypoxia (0% O₂; 100% N₂), followed by heart collection for transcriptomic and metabolomic profiling. **(B)** Time to terminal loss of coordinated cardiac electrical activity during 0% O₂ exposure, defined by sustained cessation of spontaneous respiration accompanied by progressive ECG disorganization under deep anesthesia. Bars indicate mean ± s.e.m. Exact P values are shown. **(C)** Pulmonary edema assessed by wet-to-dry lung weight ratios measured immediately following terminal hypoxic exposure (n = 3) and BMR (n = 3). Wet weight was obtained at the time of tissue excision, dry weight was measured after 24 and 48h at 60 °C. Data are presented as mean ± s.e.m. Statistical significance was determined using two-tailed unpaired t tests. Exact *P* values are shown. **(D)** Volcano plot showing differential gene expression between hypoxic BMR and hypoxic mouse hearts. Each point represents a gene plotted by log₂ fold change and −log₁₀ adjusted P value; dashed lines indicate the significance and fold-change thresholds (see Methods). **(E)** Unsupervised hierarchical clustering heatmap of significantly differentially expressed genes under hypoxia (normalized expression), showing species-specific transcriptional patterns. Rows represent genes and columns represent individual samples. **(F)** Gene set enrichment analysis (GSEA) of hypoxia-associated pathways comparing BMR versus mouse hearts under hypoxia, displayed as a dot plot of normalized enrichment score (NES). Pathways enriched in BMR are shown as “Activated” (positive NES), whereas pathways relatively depleted are shown as “Suppressed” (negative NES). Dot size indicates gene set size, and color indicates adjusted P value. **(G)** Gene-level heatmap summarizing log₂ fold-change values (BMR relative to mouse) for selected hypoxia-responsive genes grouped by functional category, including autophagy and vesicular trafficking, cell-cycle regulation, DNA damage response and repair, mitochondrial maintenance, and stress response/inflammation (hypoxia and normoxia shown as indicated). Asterisks denote significance of differential expression based on adjusted P values (see Methods). **(H)** Pathway enrichment analysis of hypoxia-responsive metabolites in BMR hearts, displayed as a dot plot (x-axis, −log₁₀[P]). Dot size reflects the number of metabolites mapped to each pathway (“Hits”), and color denotes FDR. Adjusted P values/FDR were used for differential expression and enrichment analyses as described in Methods.

In mice, hypoxic exposure induced rapid physiological deterioration characterized by early cessation of spontaneous respiration and loss of coordinated physiological activity. Based on combined respiratory and cardiac criteria, mice reached a terminal physiological endpoint within approximately ∼180 s of exposure. In contrast, BMRs exhibited a markedly delayed decline, maintaining coordinated physiological function for substantially longer durations, with terminal endpoints occurring only after ∼600 s of hypoxia. Accordingly, the duration of coordinated physiological activity was significantly prolonged in BMRs relative to mice (Fig. 1B).

Consistent with these physiological differences, hypoxia exposure resulted in pronounced pulmonary edema in mice, as indicated by significantly increased wet-to-dry lung weight ratios, whereas BMR lungs remained indistinguishable from normoxic controls (Fig. 1C). Together, these findings demonstrate that BMRs sustained cardiac and respiratory activity for extended periods under hypoxia and exhibit reduced end-organ injury compared to hypoxia-sensitive species.

### Blind mole-rats deploy a cardioprotective transcriptional program under severe hypoxia

To investigate how extreme oxygen deprivation alters cardiac gene expression across species, we performed comparative transcriptomic profiling of hearts from BMR and mice exposed to either normoxic conditions (∼21% O₂) or severe hypoxia (0% O₂; 100% N₂) (Fig. 1A).

Under normoxic conditions, global transcriptomic analyses revealed distinct species-specific expression landscapes, indicating divergent basal cardiac transcriptional states in BMR and mouse hearts (Fig. S1A,B; Data S1). Exposure to severe hypoxia further amplified these differences with clear separation of samples by species and oxygen status (Fig. 1D,E; Data S2). Differential expression analysis identified 2,539 genes significantly upregulated, and 3,055 genes significantly downregulated in hypoxic BMR hearts relative to hypoxic mouse hearts (FDR < 0.05).

To interpret the biological functions of hypoxia-responsive genes in BMR hearts, we performed pathway enrichment analysis on differentially expressed genes (Data S3). The analysis revealed a highly structured transcriptional response in BMR hearts, characterized by selective activation and suppression of distinct biological processes (Fig. 1F). Pathways related to genome maintenance including *DNA repair*, *homologous recombination*, and *chromosome organization* were among the most strongly enriched pathways. In contrast, immune- and inflammation-associated pathways, including *adaptive immune response, immune effector processes*, *cytokine-mediated signaling*, were broadly suppressed in BMR hearts (Fig. 1F; Fig. S2). These pathway-level signatures were supported by coordinated gene-level expression changes.

Consistent with enrichment analyses, hypoxic BMR hearts exhibited elevated expression of genes involved in base- and nucleotide-excision repair (e.g., *APEX1*, *XPA*), Fanconi anemia signaling (e.g., *FANCD2*), and homologous recombination (e.g., *RAD51*, *BRCA1/2*), together with upstream checkpoint regulators (Fig. 1G). Conversely, reduced expression of interferon-stimulated genes (e.g., *IFIT3*, *ISG15*), stress-activated MAPK/JNK signaling components (e.g., *MAP3K11*, *MAPK10*), and cytokine-response genes mirrored the suppression of immune-related pathways observed at the enrichment level (Fig. 1F, G), reflecting a restrained inflammatory and stress-response profile during oxygen deprivation.

In addition to genome maintenance, hypoxic BMR hearts showed enrichment of processes related to cellular organization and mitochondrial regulation, including *chromosome organization, microtubule-based movement, mitochondrial gene expression, mitochondrial translation*, and *tRNA metabolic processes* (Fig. 1F), pointing to coordinated regulation of nuclear and mitochondrial programs under hypoxic stress. Gene-level analysis further revealed coordinated expression changes in transcripts associated with mitochondrial maintenance and gene expression, as visualized in the focused heatmap (Fig. 1G), including regulators of mitochondrial transcription and proteostasis such as *TFAM*, *TFB1M*, *TFB2M*, and *YME1L1*.

These transcriptomic analyses reveal pronounced species-specific differences in cardiac gene expression at baseline and during acute hypoxia. BMR hearts display a transcriptional profile characterized by enhanced genome maintenance capacity and reduced inflammatory signalling. In parallel, enrichment of pathways related to mitochondrial regulation suggests coordinated engagement of programs supporting organelle homeostasis during oxygen deprivation.

### Blind mole-rats engage coordinated metabolic remodeling under severe hypoxia

To determine whether the transcriptional adaptations observed in BMR hearts were accompanied by corresponding metabolic changes, we performed untargeted LC–MS–based metabolomic profiling of cardiac tissue from BMRs and mice under normoxic and severe hypoxic conditions. In total, 44 metabolites were quantified under normoxic conditions, and a total of 81 metabolites were quantified under hypoxic conditions, all of which were significantly altered in BMR vs mouse hearts (FDR < 0.05) (Data S4, Data S5).

Pathway-level enrichment analysis revealed extensive metabolic reorganization in BMR hearts during hypoxia as compared to mouse (Fig. 1H; Data S5). Because these data are inherently observational, pathway enrichments are interpreted as adaptive correlations of hypoxia tolerance rather than direct causal drivers. Under normoxic conditions, pathway enrichment differences between species were limited and characterized by low pathway hit counts (Fig. S4; Data S4). In contrast, severe hypoxia elicited a robust and structured enrichment pattern in BMR hearts, indicative of an active, and coordinated metabolic response to oxygen deprivation (Fig. 1H).

Among hypoxia-responsive pathways, arginine–proline metabolism emerged as one of the most prominently enriched metabolic axes in BMR hearts (Fig. 1H). This pathway is tightly linked to mitochondrial redox regulation and amino-acid–derived stress adaptation. Notably, transcriptomics and metabolomic datasets showed concordant directional changes for selected pathways including elevated expression of *PRODH,* the rate-limiting enzyme in proline metabolism (Data S2). Additional enriched pathways included redox- and membrane-associated processes, including glutathione metabolism and lipid remodeling pathways consistent with metabolic conditions permissive for mitochondrial maintenance during hypoxic stress. Complementarily, hypoxia-enriched pathways included *alanine–aspartate–glutamate metabolism, the pentose phosphate pathway, and purine metabolism*, indicate engagement of metabolic modules that support redox buffering (e.g., NADPH generation), sustain glutathione pools, and limit oxidative stress under oxygen-limited conditions (Fig. 1H).

Polyamine-associated metabolites, notably spermidine and spermine, were elevated in hypoxic BMR hearts compared to mouse. This pattern was supported by concordant transcriptional changes consistent with increased polyamine biosynthetic capacity (eg. *Odc1, Mycbp*) and reduced polyamine turnover (eg., *Sat1, Smox*) in BMR hearts (Data S2), indicating increased polyamine availability in hypoxic BMR hearts. Given the established roles of polyamines in mitochondrial membrane stabilization, autophagy induction, and cellular stress resistance [38–43], these changes align with the broader transcriptional signatures of elevated mitochondrial quality-control and autophagy-related programs observed in BMR hearts.

Pathway analysis further identified enrichment of tricarboxylic acid (TCA) cycle–associated metabolites in hypoxic BMR hearts (Fig. 1H). Although oxygen limitation constrains oxidative TCA flux, the concurrent enrichment of *arginine-, proline-, glutamate-*, and *α-ketoglutarate–*associated pathways is biochemically consistent with amino acid–derived anaplerotic inputs that sustain TCA intermediates during hypoxia rather than increased oxidative metabolism.

Collectively, these data indicate that BMR hearts undergo coordinated metabolic remodeling during severe hypoxia, with pathway-level changes that align with the transcriptomic signatures observed in parallel and contrast with the more limited metabolic response in hypoxia-sensitive mouse hearts.

### Blind mole-rats suppress mitochondrial respiration and RET-derived ROS under acute hypoxia

To determine whether the metabolic rewiring observed in BMR hearts was accompanied by intrinsic mitochondrial adaptations, we assessed mitochondrial respiratory function and ROS generation in isolated cardiac mitochondria. High-resolution respirometry revealed marked species-specific differences in respiratory state engagement during oxygen deprivation.

In mouse mitochondria, both Complex I–linked (pyruvate/malate) and Complex II–linked (succinate) respiration remained unchanged between normoxic and hypoxic conditions across OXPHOS and ET states (Fig.2 A,C). These observations indicate that mouse mitochondria do not adjust respiratory engagement when oxygen availability collapses. By contrast, BMR mitochondria displayed a coordinated and robust downregulation of respiratory activity under hypoxia. Both complex I– and complex II–supported OXPHOS, as well as maximal ET capacity, were significantly reduced in hypoxic BMR samples relative to normoxia (Fig. 2B,D). This broad suppression across respiratory flux indicates an intrinsic mitochondrial response that limits electron flow through the electron transport chain when oxygen is scarce, minimizing electron pressure under conditions where oxygen is unavailable as a terminal electron acceptor.

**Figure 2.**
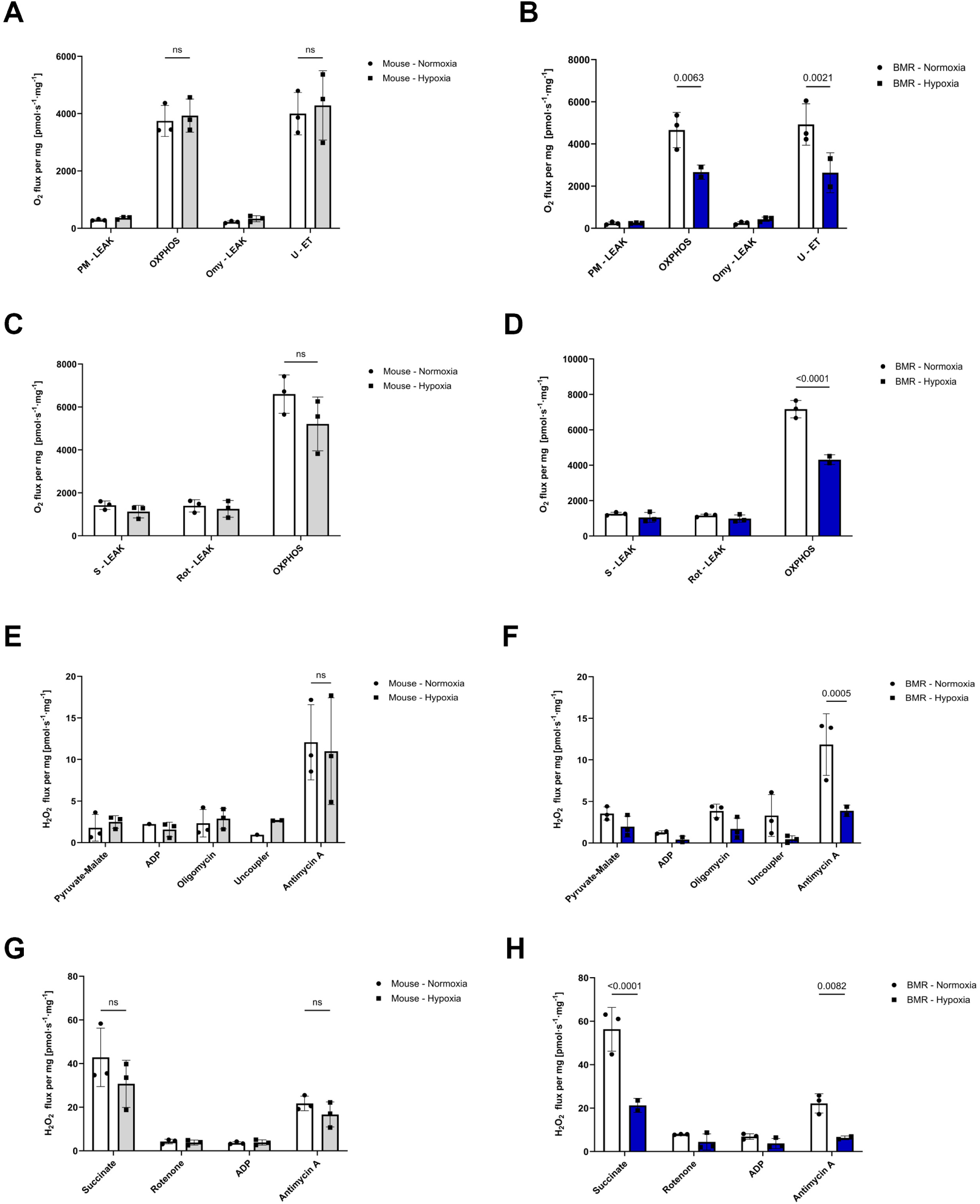
Hypoxia suppresses mitochondrial respiration and RET-associated ROS production in BMR hearts. Oxygen consumption rates (O₂ flux per mg protein) and hydrogen peroxide production (H₂O₂ flux per mg protein) were measured in isolated heart mitochondria from mouse (left panels) and blind mole-rat (BMR; right panels) following normoxic or hypoxic exposure (n = 3 per group). Mitochondrial respiration was assessed using sequential substrate–uncoupler–inhibitor titration protocols in a high-resolution respirometry system. **(A, B)** Complex I–linked respiration supported by pyruvate/malate during oxidative phosphorylation (OXPHOS) and maximal electron transfer (ET) capacity in mouse (A) and BMR (B) mitochondria. **(C, D)** Complex II–linked respiration supported by succinate during OXPHOS and ET capacity in mouse (C) and BMR (D) mitochondria. **(E, F)** H₂O₂ emission during forward electron transport (FET) supported by pyruvate/malate in mouse (E) and BMR (F) mitochondria. **(G, H)** Reverse electron transport (RET)–associated H₂O₂ production induced by succinate in the absence of rotenone in mouse (G) and BMR (H) mitochondria. Data are presented as mean ± s.e.m. Statistical significance was determined using two-tailed unpaired t tests. Exact P values are shown.

To assess whether these respiratory changes were accompanied by altered mitochondrial ROS generation during forward electron transport (FET), we quantified hydrogen peroxide (H₂O₂) emission across sequential respiratory states supported by pyruvate/malate. In mouse mitochondria, H₂O₂ release during FET remained comparable between normoxic and hypoxic conditions across all measured states (Fig. 2E), consistent with maintained respiratory activity. By contrast, BMR mitochondria exhibited a significant reduction in H₂O₂ production following hypoxic exposure, particularly under conditions associated with high electron flux through the respiratory chain (Fig. 2F). These findings indicate that hypoxia suppresses FET-associated ROS generation in BMR mitochondria.

Since reverse electron transport (RET) is a major source of pathological mitochondrial ROS, we next examined succinate-driven H₂O₂ production in the absence of rotenone. In mouse mitochondria, RET-associated H₂O₂ emission remained unchanged between normoxic and hypoxic conditions (Fig. 2G), consistent with sustained electron pressure and the lack of respiratory adjustment. In contrast, BMR mitochondria displayed a marked reduction in succinate-supported H₂O₂ release following hypoxia (Fig. 2H), accompanied by diminished sensitivity to RET-associated ROS generation. This reduction is consistent with reduced electron backflow at complex I, a key mechanism for preventing ROS overproduction when electrons accumulate upstream during oxygen scarcity.

These differences reinforce that BMR cardiac mitochondria actively suppress both FET and RET–associated ROS production during acute hypoxia, in parallel with reduced engagement of oxygen-intensive respiratory states. This intrinsic mitochondrial phenotype provides a mechanistic basis for limiting oxidative stress under extreme oxygen deprivation and aligns with the transcriptional and metabolic adaptations observed *in vivo*.

### Acute hypoxia selectively activates AMPK–mTOR–ULK1–dependent mitophagy in Blind mole-rats

Given the mitochondrial restraint observed in hypoxic BMR hearts, we next sought to identify regulatory mechanisms that could engage in parallel to this phenotype to preserve organelle integrity. Unbiased transcriptomic analyses captured this at systems level, revealing coordinated expression changes in transcripts associated with autophagosome biogenesis and vesicular trafficking (Fig. 1G).

BMR hearts exhibited elevated expression of genes involved in early phagophore assembly, including *RB1CC1/FIP200*, *PIK3R4/VPS15*, and *WIPI2*, together with the selective autophagy adaptor *TAX1BP1* (Fig. 1G; Data S1; Data S2) under normoxic and hypoxic conditions. During hypoxia, this primed state was accompanied by broader transcriptional reinforcement of the autophagy machinery, including induction of genes involved in autophagosome elongation and ATG8 processing (e.g., *ATG12*, *ATG4C*, *ATG4D*), as well as upregulation of vesicular trafficking components linked to autophagosome maturation and cargo handling (Fig. 1G). Protein-protein interaction and gene ontology analyses revealed enrichment of modules associated with *autophagy*, *organelle disassembly*, and *autophagy of mitochondrion* in BMR hearts relative to mouse under both normoxic and hypoxic conditions (Fig. S3A,B; Fig. 3A,B).

**Figure 3.**
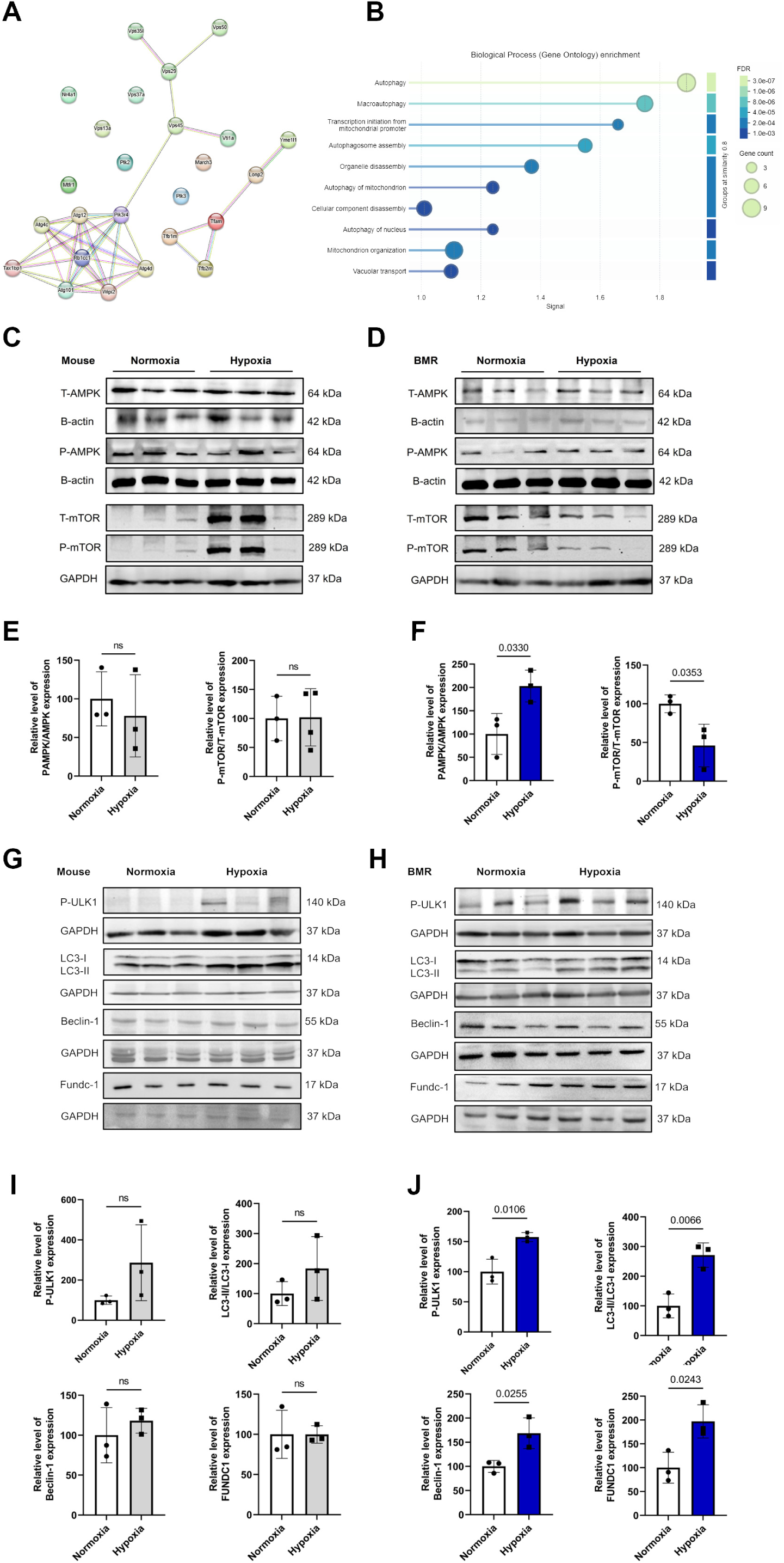
Hypoxia engages AMPK–mTOR–ULK1–dependent mitophagy in BMR hearts. **(A)** Protein–protein interaction network constructed from genes significantly upregulated in blind mole rat (BMR) hearts under hypoxic conditions. Nodes represent individual proteins, and edges denote known or predicted functional interactions. **(B)** Gene Ontology (GO) biological process enrichment analysis of hypoxia-responsive transcripts in BMR hearts. Enriched terms are displayed as a bubble plot, with bubble size proportional to the number of genes contributing to each term and color indicating false discovery rate (FDR). Prominently enriched processes include autophagy, macroautophagy, mitophagy, autophagosome assembly, mitochondrial organization, and organelle disassembly. **(C, D)** Representative immunoblots of total and phosphorylated AMPK and mTOR in mouse (A) and blind mole-rat (BMR; B) hearts under normoxia and severe hypoxia (0% O₂). β-actin was used as a loading control for AMPK blots, and GAPDH was used as a loading control where indicated. **(E, F)** Quantification of p-AMPK/AMPK and p-mTOR/mTOR ratios in mouse (C) and BMR (D) hearts. **(G, H)** Representative immunoblots of ULK1 phosphorylation and autophagy/mitophagy-related proteins in mouse (E) and BMR (F) hearts under normoxia and hypoxia, including phosphorylated ULK1 (p-ULK1), LC3 (LC3-I and LC3-II), Beclin-1, FUNDC1, and cleaved caspase-3. GAPDH was used as a loading control. **(I, J)** Quantification of p-ULK1, LC3-II/LC3-I ratio, Beclin-1, and FUNDC1 expression in mouse (G) and BMR (H) hearts, normalized to GAPDH as indicated. Data are presented as mean ± s.e.m.; n = 3 biological replicates per condition. Statistical significance was determined using two-tailed unpaired t tests. Exact P values are shown.

However, transcripts encoding upstream regulators of autophagy initiation were not significantly altered under hypoxia. As autophagy initiation is primarily regulated at the post-translational level, we next evaluated activation of upstream autophagy regulators at the protein level. In particular, we examined the AMPK–mTOR–ULK1 signaling axis, a central pathway linking energetic stress to activation of the ULK1 autophagy initiation complex. Immunoblot analyses revealed robust activation of AMPK signaling in BMR hearts exposed to 0% O₂, reflected by a significant increase in the p-AMPK/AMPK ratio compared with normoxic controls (Fig. 3A–D). In parallel, mTOR activity was reduced, as indicated by a decreased p-mTOR/mTOR ratio (Fig. 3D). Concomitantly, pULK1 abundance was increased in hypoxic BMR hearts (Fig. 3E–F), consistent with activation of the autophagy initiation machinery.

These signaling changes were accompanied by increased levels of mitophagy-associated proteins, including FUNDC1 and Beclin-1, together with elevated LC3-II levels and a higher LC3-II/LC3-I ratio (Fig. 3H; Fig. S5B,D), consistent with enhanced autophagosome formation and engagement of the autophagy machinery [29]. Levels of cleaved caspase-3 were reduced in hypoxic BMR hearts (Fig. S5F,H), consistent with attenuation of apoptotic signaling.

In contrast, mouse hearts exposed to the same hypoxic challenge did not exhibit significant changes in AMPK or mTOR activation status, nor in ULK1 phosphorylation or the abundance of the mitophagy-associated proteins examined (Fig. 3G). Although LC3-II levels increased in hypoxic mouse hearts (Fig. S5A,C), the LC3-II/LC3-I ratio remained unchanged (Fig. 3G). Hypoxic mouse hearts also displayed increased cleaved caspase-3 (Fig. S5E, G), consistent with activation of apoptotic signaling in the absence of coordinated mitophagy induction.

Together, these data indicate that BMR hearts combine a transcriptionally primed autophagy machinery with hypoxia-induced post-translational activation of the AMPK–mTOR–ULK1 axis, enabling coordinated engagement of mitophagy under severe oxygen deprivation.

### Mitophagy preserves cardiomyocyte survival and mitochondrial integrity in blind mole-rats during hypoxia–reoxygenation

To determine whether mitophagy is functionally required for hypoxia tolerance in BMRs, we isolated primary cardiomyocytes from adult male BMRs and rats and subjected them to hypoxia–reoxygenation (H/R; 30 min hypoxia followed by 2 h reoxygenation) (Fig. 4A). Under H/R conditions, BMR cardiomyocytes preserved viability at levels comparable to normoxic controls, indicating cell-intrinsic resistance to hypoxic injury. In contrast, rat cardiomyocytes exhibited a significant reduction in viability relative to normoxia (Fig. 4B).

**Figure 4.**
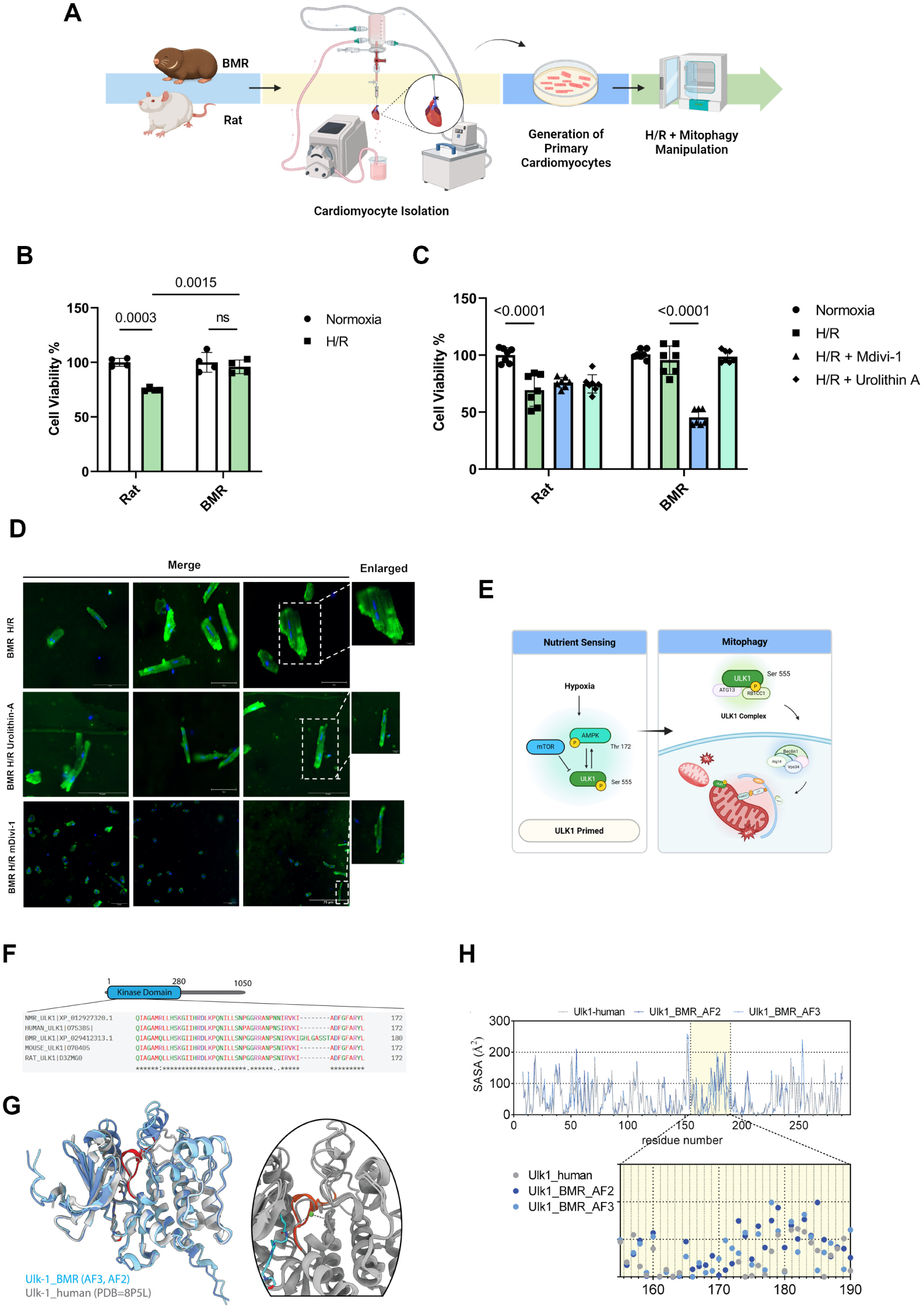
Mitophagy is required for hypoxia–reoxygenation resistance in BMR cardiomyocytes and is linked to a BMR-specific ULK1 variant. **(A)** Schematic overview of the experimental workflow. Primary cardiomyocytes isolated from blind mole-rats (BMRs) and rats were subjected to hypoxia–reoxygenation (H/R; 30 min hypoxia followed by 2 h reoxygenation) and treated with pharmacological modulators of mitophagy prior to assessment of cell viability and mitochondrial morphology. **(B)** Cell viability of rat and BMR cardiomyocytes under normoxic conditions and following H/R. **(C)** Effects of mitophagy modulation on cardiomyocyte viability following H/R. Cells were treated with the mitophagy inhibitor Mdivi-1 (20 μM) or the mitophagy activator urolithin A (5 μM) during H/R. **(D)** Representative fluorescence micrographs of BMR cardiomyocytes stained with MitoTracker Green (mitochondria) and Hoechst 33342 (nuclei) under the indicated conditions. Images were acquired at 40X magnification; representative images are shown from at least three independent biological replicates per condition. **(E)** Conceptual schematic illustrating hypoxia-induced AMPK–mTOR–ULK1 signaling and priming of the mitophagy machinery, and downstream execution of mitochondrial clearance via the ULK1 complex. **(F)** Multiple sequence alignment of ULK1 orthologs from human, mouse, rat, naked mole-rat (NMR), and BMR. An eight–amino-acid insertion (residues 164–171) unique to BMR ULK1 is located within a highly conserved region of the kinase domain. **(G)** Structural mapping of the BMR-specific ULK1 insertion. The kinase domain of human ULK1 (PDB: 8P5L) was used as a reference structure. Predicted BMR ULK1 kinase-domain structures generated by AlphaFold2 and AlphaFold3 are shown, with the insertion highlighted relative to the catalytic core. **(H)** Solvent-accessible surface area (SASA) profiles of human ULK1 (experimental structure) and BMR ULK1 (AlphaFold2 and AlphaFold3 predictions). The inset highlights the 164–171 insertion, demonstrating altered surface exposure and local topology in the BMR variant. Data in (B) and (C) are presented as mean ± s.e.m.; n = 3 independent biological replicates per condition. Statistical significance was determined using two-way ANOVA with multiple-comparisons correction. Exact P values are shown.

To directly assess the contribution of mitophagy to this phenotype, we pharmacologically modulated mitophagy during H/R. Modulation with the mitophagy inhibitor Mdivi-1 significantly reduced the viability of BMR cardiomyocytes under H/R compared with untreated H/R samples, whereas treatment with the mitophagy activator urolithin A preserved viability at levels comparable to normoxic conditions (Fig. 4C). In contrast, rat cardiomyocytes displayed markedly reduced viability following H/R, and neither mitophagy inhibition nor activation significantly altered survival, indicating that mitophagy-dependent protection is selectively engaged in BMR cardiomyocytes during hypoxic stress.

To determine whether mitophagy modulation also influences mitochondrial structural integrity during H/R, we visualized mitochondrial morphology in BMR cardiomyocytes using MitoTracker Green staining. Under normoxic conditions, BMR cardiomyocytes displayed a dense, reticular mitochondrial network with elongated and aligned mitochondria characteristics of cardiac cells. Following H/R, mitochondrial morphology in untreated BMR cardiomyocytes remained largely preserved, consistent with their maintained viability (Fig. 4D). In contrast, pharmacological perturbation of mitophagy with Mdivi-1 resulted in mitochondrial rounding, indicative of disrupted mitochondrial architecture. Activation of mitophagy with urolithin A maintained mitochondrial morphology comparable to normoxic controls (Fig. 4D).

Together, these results demonstrate that mitophagy is functionally required to preserve both cardiomyocyte survival and mitochondrial integrity in BMRs during H/R stress. This selective reliance on mitophagy in BMR cardiomyocytes prompted us to examine upstream molecular components involved in mitophagy initiation. The AMPK–mTOR–ULK1 axis represents a conserved signaling module that regulates mitophagy through ULK1 activation (Fig. 4E), leading us to assess sequence conservation of its core components across species.

### A blind mole-rat–specific ULK1 variant enhances survival of rat cardiomyocytes during hypoxia–reoxygenation

To identify lineage-specific features that may contribute to mitophagy-associated hypoxia resilience in BMRs, we examined sequence conservation of core components of the AMPK–mTOR–ULK1 pathway across species and identified a BMR-specific insertion within the ULK1 kinase domain (Fig. 4F). Multiple sequence alignment of ULK1 orthologs from human, mouse, rat, naked mole-rat, and BMR revealed an eight–amino-acid insertion (residues 164–171) uniquely present in BMR ULK1 within the kinase domain (Fig. 4F).

To place this insertion in structural context, we modeled the BMR ULK1 kinase domain using AlphaFold2 and AlphaFold3 and aligned the predicted structures to the human ULK1 crystal structure (PDB ID: 8p5l), which was co-crystallized with a kinase inhibitor (Fig. 4G). The predicted BMR models exhibited global pLDDT scores of 83 and 86, respectively, and moderate backbone deviations from the human structure (RMSD 5.29 Å for AF2; 5.27 Å for AF3), with the largest differences localized to the loop containing the BMR-specific insertion. Total solvent-accessible surface area (SASA) was higher in the BMR models than in the human kinase domain (154 and 159 nm² versus 135 nm²), while per-residue SASA profiles were closely matched outside the insertion region. In the vicinity of residues 164–171, the BMR models displayed increased SASA and altered local surface topology compared with human ULK1 (Fig. 4H).

Notably, the BMR-specific insertion is positioned within an extended loop adjacent to the catalytic site and ATP-binding pocket in the reference human structure and contains two serine residues and one threonine residue, consistent with the presence of potential regulatory phosphorylation sites. This region does not overlap with reported ULK1 protein–protein interaction interfaces but contributes to a more exposed surface patch near the active site in the BMR models.

We next examined whether the BMR-specific ULK1 insertion confers hypoxia tolerance in a heterologous system. To assess its functional impact, we introduced the insertion into H9c2 rat cardiomyocytes using CRISPR–Cas9–mediated knock-in at the endogenous ULK1 locus (Fig. 5A). Precise genomic integration was confirmed by Sanger sequencing (Fig. S6, S7; Table S2).

**Figure 5.**
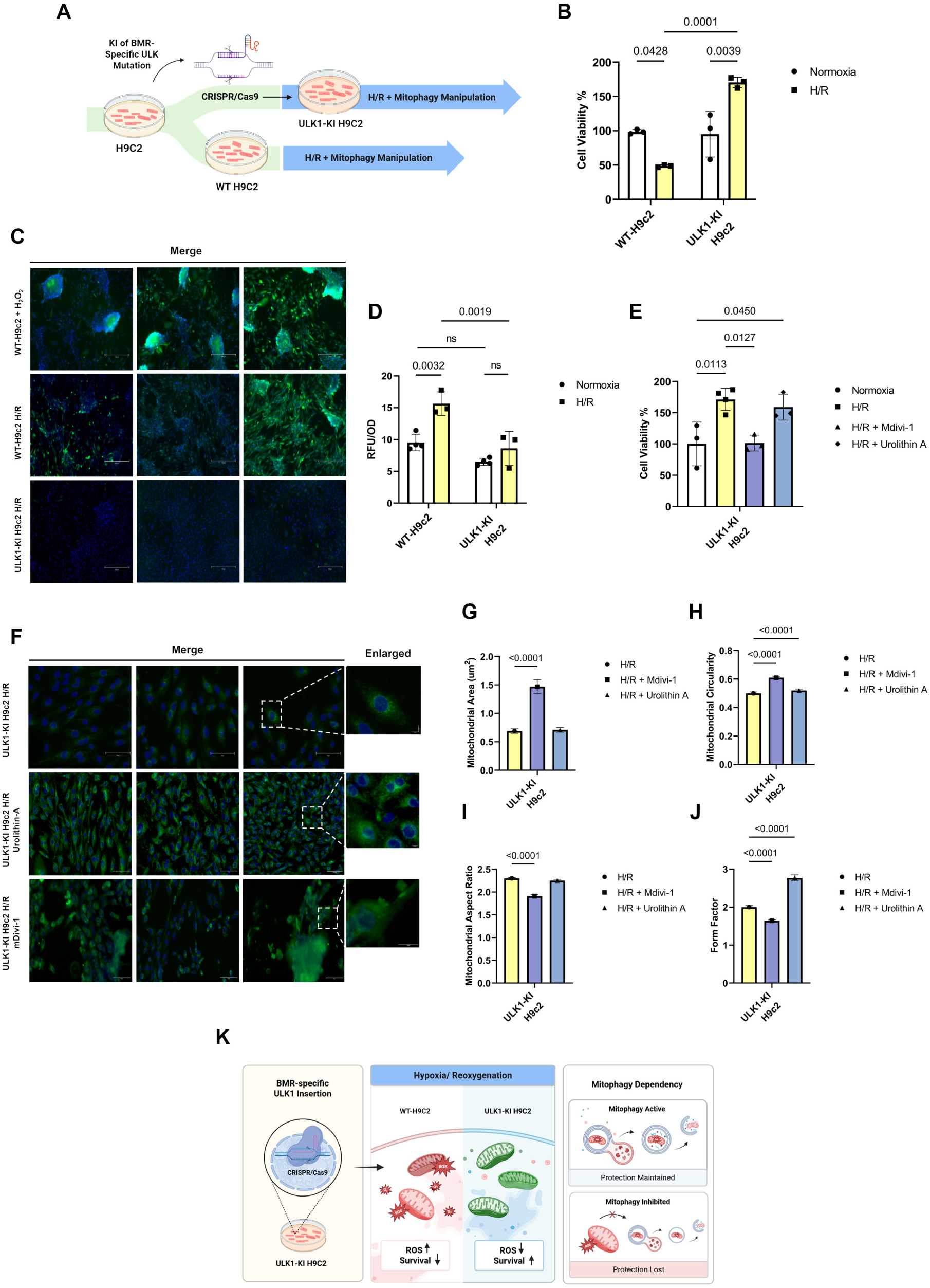
A BMR-specific ULK1 insertion confers mitophagy-dependent tolerance to hypoxia–reoxygenation stress in rat cardiomyocytes. **(A)** Schematic overview of CRISPR–Cas9–mediated knock-in (KI) of the BMR-derived ULK1 insertion into H9c2 rat cardiomyocytes. Wild-type (WT) and ULK1-KI cells were subjected to hypoxia–reoxygenation (H/R) with or without pharmacological modulation of mitophagy prior to downstream analyses. **(B)** Cell viability of WT and ULK1-KI cells under normoxia and following H/R (24 h hypoxia followed by 24 h reoxygenation). **(C)** Representative fluorescence microscopy images showing intracellular reactive oxygen species (ROS) levels detected by H₂DCFDA staining in WT and ULK1-KI cells following H/R. H₂O₂-treated cells (100 µM) are shown as a positive control. Images were acquired at 10X magnification; representative images are shown from at least three independent biological replicates per condition. **(D)** Quantification of intracellular ROS levels measured by H₂DCFDA fluorescence and normalized to cell viability assessed by CCK-8 in WT and ULK1-KI cells under normoxic and H/R conditions. ROS values were normalized to the WT normoxia group. **(E)** Effects of mitophagy modulation on cell viability following H/R in ULK1-KI cells. Cells were treated with urolithin A (5 µM), Mdivi-1 (20 µM), or vehicle control during H/R. **(F)** Representative fluorescence micrographs of ULK1-KI cells stained with MitoTracker Green (mitochondria) and Hoechst 33342 (nuclei) under the indicated conditions. Images were initially acquired using a 20X objective. Selected regions of interest (ROIs) were digitally magnified (3X) using Fiji/ImageJ software to visualize mitochondrial details. A scale bar of 75 µm was applied to the representative images. **(G–J)** Quantitative morphometric analysis of mitochondrial networks in ULK1-KI cells following H/R, including mitochondrial area (G), circularity (H), aspect ratio (I), and form factor (J), under vehicle, Mdivi-1, or urolithin A treatment. **(K)** Conceptual model summarizing the effects of the BMR-specific ULK1 insertion on mitophagy-dependent protection during H/R. ULK1 knock-in promotes mitochondrial quality control, limits ROS accumulation, and preserves cell survival under H/R stress, whereas inhibition of mitophagy abrogates this protective phenotype. Data are presented as mean ± s.e.m. unless otherwise indicated. For viability and morphometric analyses, n = 3 independent biological replicates per condition. Statistical significance was determined using two-way ANOVA with multiple-comparisons correction or unpaired two-tailed t tests, as indicated. Exact P values are shown.

Wild-type (WT) and ULK1 knock-in (KI) H9c2 cells were then subjected to H/R (H/R; 24 h hypoxia followed by 24 h reoxygenation) or maintained under normoxia. WT cells displayed a marked reduction in viability following H/R compared with normoxic controls (Fig. 5B). In contrast, ULK1 KI cells retained significantly greater viability under H/R, demonstrating that the BMR-specific insertion is sufficient to confer a survival advantage in a hypoxia-intolerant cellular background (Fig. 5B; Fig. S8).

### A Blind mole-rat–specific ULK1 variant attenuates oxidative stress during hypoxia–reoxygenation

Given the enhanced survival of ULK1 KI cells under H/R, we next assessed whether this phenotype was associated with altered intracellular oxidative stress. ROS levels were measured in WT and ULK1 KI cells using the H₂DCFDA assay. To account for potential differences in cytotoxicity, fluorescence signals were normalized to cell viability as assessed by CCK-8.

Representative fluorescence images revealed a pronounced increase in ROS accumulation in WT cells following H/R, whereas ULK1 KI cells displayed markedly reduced ROS signals (Fig. 5C). Quantitative analysis confirmed that ROS levels in ULK1 KI cells following H/R were significantly lower than those observed in WT cells and were not significantly different from normoxic baselines (Fig. 5D; Fig. S9). Together, these results indicate that introduction of the BMR-specific ULK1 variant is associated with reduced intracellular ROS accumulation during hypoxia–reoxygenation stress in rat cardiomyocytes.

### ULK1-dependent mitophagy is required for mitochondrial integrity and survival in knock-in cardiomyocytes

To determine whether the survival advantage conferred by ULK1 knock-in depends on intact mitophagy signaling, we pharmacologically modulated mitophagy during H/R. In WT cells, inhibition of mitophagy with Mdivi-1 did not significantly alter viability relative to untreated H/R controls (Fig. 5B). In contrast, in ULK1 KI cells, Mdivi-1 treatment significantly reduced viability following H/R, whereas activation of mitophagy with urolithin A preserved cell viability (Fig. 5E). These results indicate that the protective effect associated with the BMR-specific ULK1 variant depends on intact mitophagy signaling.

We next examined whether mitophagy modulation influenced mitochondrial organization in the ULK1 KI system. Following H/R, untreated ULK1 KI cells retained an elongated, filamentous mitochondrial network comparable to normoxic controls (Fig. 5F). Pharmacological perturbation of mitophagy using Mdivi-1 resulted in pronounced mitochondrial fragmentation and rounding, whereas urolithin A treatment preserved a reticular, elongated mitochondrial architecture (Fig. 5F; Fig. S10). Quantitative morphometric analysis revealed that Mdivi-1 treatment increased mitochondrial area and circularity while reducing aspect ratio, consistent with mitochondrial swelling and loss of network integrity (Fig. 5G). In contrast, urolithin A maintained mitochondrial area closer to control levels and increased form factor, indicative of greater network complexity and branching (Fig. 5G).

Collectively, these findings demonstrate that introduction of the BMR-specific ULK1 insertion into hypoxia-intolerant rat cardiomyocytes is sufficient to enhance tolerance to H/R stress, characterized by preserved cell viability, reduced oxidative stress, and maintenance of mitochondrial network integrity. This protective phenotype is dependent on intact mitophagy signaling and is lost upon pharmacological perturbation of mitophagy (Fig. 6K).

## Discussion

Mammalian hearts are highly sensitive to oxygen limitation, yet BMRs thrive in chronically hypoxic subterranean environments and maintain cardiovascular function under sustained low-oxygen conditions [10]. Here, we define a cardioprotective program in BMRs that preserves mitochondrial maintenance through coordinated transcriptional, metabolic, and mitochondrial adaptations. Central to this cardiac resilience is AMPK–mTOR–ULK1 mediated mitophagy which links energetic stress to mitochondrial turnover, offering a coherent strategy to sustain organelle function under extreme hypoxia.

At the organismal level, BMRs survived significantly longer and maintained pulmonary stability during acute 0% O₂ exposure compared to mouse. This physiological tolerance was mirrored by a distinct molecular response. Under hypoxia, BMR hearts engaged genome-maintenance programs while displaying restrained induction of interferon- and stress-associated signaling pathways, consistent with reduced activation of damage-responsive and inflammatory pathways. This attenuated transcriptional response contrasts with the canonical hypoxic response observed in hypoxia-sensitive species and supports downstream metabolic and mitochondrial adaptations.

At the mitochondrial level, this hypoxia-tolerant phenotype is accompanied by intrinsic suppression of oxygen-intensive respiratory states. Direct measurements of mitochondrial function revealed reduced oxidative phosphorylation and electron transfer capacity in hypoxic BMR mitochondria, a configuration expected to lower electron pressure within the respiratory chain and thereby limit ROS generation. Consistent with this model, RET-associated H₂O₂ emission, a major source of mitochondrial ROS during ischemic stress [30] was markedly suppressed in BMR mitochondria but not in mouse. By constraining respiratory flux and RET, BMR hearts limit mitochondrial ROS production and protect organelle integrity during oxygen deprivation.

Metabolomic profiling provides supportive context for this mitochondrial phenotype. Under hypoxia, BMR hearts show enrichment of amino-acid–linked, redox-associated, and lipid-related pathways with a metabolic state that supports redox buffering and membrane maintenance [31–38]. A second notable feature was the increased abundance of polyamines, including spermidine and spermine, in hypoxic BMR hearts. Given prior reports linking polyamines to stress resilience and mitochondrial membrane stability in cardiac models [40–45], this enrichment aligns with a metabolic environment that may further support mitochondrial integrity during oxygen limitation.

Upstream, hypoxia engaged the AMPK–mTOR–ULK1 axis in BMR hearts, coupling energetic stress to mitophagy activation [25–27]. Notably, autophagy- and mitochondrial organization–related pathways were enriched at the transcript level even under normoxia consistent with constitutive expression of components required for mitochondrial surveillance. In contrast, hypoxia-dependent activation occurred predominantly through post-translational modification of core regulators. This layered organization of transcriptional priming coupled to post-translational activation may enable rapid swift and efficient mitochondrial turnover in response to acute oxygen deprivation. Consistent with this signaling profile, primary BMR cardiomyocytes maintained viability and mitochondrial network organization during H/R, and pharmacological perturbation of mitophagy abolished this resilience. These results support that mitophagy is not merely elevated at the transcript or protein level but is functionally required for cardiomyocyte survival during hypoxic stress.

A key mechanistic insight is the identification of an eight–amino acid insertion in the BMR ULK1 kinase domain that is absent from other examined mammals. Structural modelling positions this insertion in a flexible loop adjacent to the catalytic region, suggesting altered accessibility or regulatory interactions without disrupting global kinase architecture. Functionally, introduction of this insertion into rat cardiomyocytes enhanced survival during H/R, accompanied by reduced ROS accumulation and preservation of mitochondrial network organization, indicating that BMR-specific ULK1 variant is sufficient to confer core features of hypoxia-tolerant phenotype observed in native BMR cardiomyocytes.

Notably, this protection is sensitive to intact mitophagy signaling. In ULK1 KI cells, mitophagy inhibition abrogated the survival advantage and promoted mitochondrial fragmentation together with loss of network integrity, whereas mitophagy activation preserved mitochondrial architecture and viability. Together, these data establish that ULK1 insertion enhances cellular resilience through a mitophagy-dependent mechanism that preserves mitochondrial integrity and limits oxidative stress during H/R.

Collectively, our data supports a model in which BMR cardiac hypoxia tolerance in the BMR is centered on mitochondrial maintenance control that sustain function under extreme oxygen deprivation. Constitutive genome protection limits secondary stress damage; metabolic adaptations help sustain redox balance and membrane integrity; mitochondria restrict high-flux respiratory states to minimize ROS generation; and the AMPK–mTOR–ULK1 mediated mitophagy axis ensures efficient turnover of damaged mitochondria. The BMR-specific ULK1 insertion functionally augments this framework, representing a rare example of evolutionary modification of a core autophagy regulator that confers measurable resilience in a hypoxia-intolerant cellular background.

By elucidating how a naturally hypoxia-tolerant mammal integrates metabolic flexibility with mitochondrial surveillance, this study expands our understanding of endogenous cardioprotective strategies. Given the central roles of mitochondrial dysfunction, redox imbalance, and impaired mitophagy in ischemic heart disease and aging, the BMR offers a comparative framework for identifying principles that may be leveraged to enhance mitochondrial resilience and limit hypoxia-induced damage in hypoxia-intolerant systems.

## Limitation of the Study

This study has several limitations. First, experimental work in blind mole-rats (BMRs) is inherently constrained by the limited availability of animals and the technical challenges associated with long-term housing and in vivo manipulation of this subterranean species. These factors restrict sample sizes and limit the types of physiological experiments that can be performed in vivo. Second, mechanistic studies in BMRs are further limited by the absence of established blind mole-rat cell lines and genetic tools. To address this, we isolated and cultured BMR cardiomyocytes to enable cell-intrinsic functional analyses. However, isolation of viable primary cardiomyocytes from BMRs is technically challenging, yields are limited, and prolonged culture or extensive genetic manipulation is not feasible, restricting the scope of cellular perturbations that can be performed directly in this species. Third, in vivo hypoxia experiments were necessarily conducted using acute, experimentally defined exposure paradigms to permit controlled physiological monitoring and tissue collection. Although these conditions robustly revealed interspecies differences in hypoxia tolerance, they do not fully capture the chronic and fluctuating hypoxic environments experienced by blind mole-rats in their natural subterranean habitats.

## Materials and Methods

### Field collection and Animal Handling

Blind mole-rats (BMRs; *Nannospalax* spp.) were captured in the Bolu–Gerede region of Türkiye as part of a long-term field monitoring program, using standard species-specific trapping techniques widely applied in studies of subterranean rodents. Comparable field collection and handling procedures have been routinely used in previous biological and ecological investigations of *Spalax/Nannospalax* populations [46]. Briefly, young adult animals (1–6 months) were briefly removed from their nests, individually barcoded, and biometrically recorded (age, weight, sex, health status) before returning to their natal burrows. Following capture or acquisition, BMRs were housed under controlled environmental conditions adapted to their fossorial physiology, at 22 ± 2 °C and 50 ± 10% relative humidity, with ad libitum access to food and water.

Laboratory BALB/c mouse were housed under specific pathogen-free conditions in individually ventilated cages with a 12 h light/dark cycle, at an ambient temperature of 23 ± 1 °C and relative humidity of 45–65%, with ad libitum access to food and water. Adult male BMRs (9-12 months) and BALB/c mice (3-6 months) were used (n = 6 per species per condition unless otherwise indicated).

All procedures were approved by the Acıbadem University Animal Ethics Committee (ACU-HADYEK 2022/63) and performed in accordance with institutional and national guidelines.

### *In Vivo* Hypoxia Exposure

Prior to experimental procedures, animals were anesthetized by intraperitoneal injection of ketamine (90 mg/kg) and xylazine (10 mg/kg). Adequate depth of anesthesia was confirmed by the absence of pedal withdrawal and corneal reflexes before assignment to normoxic or hypoxic conditions.

For hypoxia exposure, anesthetized animals were placed in a transparent hypoxia chamber that was rapidly flushed with 100% nitrogen (N₂). Nitrogen was delivered at a pressure of 0.5 bar, resulting in complete chamber gas exchange within approximately 5–10 s, as verified by continuous gas outflow through a water-sealed outlet. Animals were exposed to hypoxic conditions for empirically determined durations selected to encompass terminal physiological failure for each species (BALB/c mice, 5 min; blind mole-rats, 10 min).

Throughout hypoxic exposure, physiological status was continuously monitored by synchronized video recording and electrocardiography (ECG). Cardiac electrical activity was recorded in real time using subcutaneous electrodes connected to a BIOPAC MP36 data acquisition system, while respiratory activity was assessed visually from video recordings. A terminal cardiorespiratory endpoint was defined by sustained cessation of spontaneous respiration accompanied by progressive disorganization and loss of coordinated cardiac electrical activity under deep anesthesia. ECG-derived heart rate signals that persisted transiently beyond this endpoint were interpreted as terminal electrical activity rather than effective cardiac function. Upon confirmation of the terminal cardiorespiratory endpoint, animals were euthanized by cervical dislocation under deep anesthesia. Hearts were immediately excised, rinsed in ice-cold phosphate-buffered saline (PBS), snap-frozen in liquid nitrogen, and stored at −80 °C until further analysis.

Pulmonary edema following normoxic or hypoxic exposure was quantified using the wet-to-dry (W/D) lung weight ratio method. At the experimental endpoint, animals were euthanized under deep anesthesia, lungs were carefully excised and weighed to obtain wet weight. Lung tissues were then dried in a ventilated oven at 60 °C for 48 h until constant mass was achieved and reweighed to determine dry weight. The W/D ratio was calculated as an index of pulmonary edema. Analyses were performed using lung tissue from *n* = 3 animals per species per condition.

### RNA Sequencing

Total RNA was isolated from snap-frozen heart tissue using the QIAGEN RNeasy Mini Kit (cat# 74104), following the manufacturer’s silica-membrane spin-column protocol. Frozen tissue (≤30 mg per sample) was handled on ice and homogenized in Buffer RLT supplemented with β-mercaptoethanol using a rotor–stator homogenizer. RNA was purified according to the manufacturer’s instructions, eluted in RNase-free water, and stored at −80 °C until further analysis. The RNA samples were examined for quality before undergoing library preparation using the Illumina TruSeq Stranded Total RNA Prep with Ribo-Zero Plus protocol (Illumina, San Diego, USA) by Tufts Genomics core facility. Single-end RNA-sequencing was performed on the Illumina NovaSeq X Plus platform at a read length of 100 bp and a depth of 33 million reads per sample.

RNA-Seq data were processed using nf-core/rnaseq (v3.18.0) [47] on Nextflow (v24.10.5) [56]. Raw reads underwent quality assessment with FastQC and were summarized in MultiQC; adapters were removed using Trim Galore (Cutadapt) [57, 58]. Quantification was performed with Salmon (v1.10.3) using selective alignment against Ensembl references; gene-level counts were aggregated via tximport/tximeta to preserve reference provenance [59, 60]. Additional QC metrics were obtained using Qualimap and RSeQC, with PCR duplication assessed by dupRadar [61–63]. Where alignments were generated (e.g., for genome indices or auxiliary metrics), STAR [64] and SAMtools [68] were used; read assignment via featureCounts (Subread) [65], duplicate marking via Picard [66], transcriptome reconstruction with StringTie [67], and coverage utilities via BEDTools [69] were employed as needed.

### Ortholog Mapping and Normalization

Gene IDs of Mouse and BMR were harmonized by identifying 1:1 orthologs using reciprocal best-hit BLASTP searches (BLOSUM45 matrix; minimum identity 30%; cutoff length 100 amino acids). Protein IDs were mapped to Ensembl gene IDs using species-specific FASTA headers. Only genes present in both species and matching top BLAST hits in both directions were retained. Counts for orthologous genes were combined into a unified matrix, and low-abundance genes (total counts ≤ 20) were filtered out. To correct for gene-length bias in interspecies comparisons, exon-sum lengths were computed from Ensembl GTF annotations. A normalization matrix was constructed and row-centered by geometric mean, then passed to DESeq2 for custom size-factor estimation.

### Differential Expression Analysis

Differential expression analysis was performed using DESeq2 (v1.46.0) [48] in R (v4.4.2) with design ∼ Group. Pairwise contrasts were computed between Mouse and BMR under Normoxia and Hypoxia conditions. Statistical significance was assessed using the Wald test with Benjamini–Hochberg correction. Variance-stabilizing transformation (VST) was applied for visualization. Principal component analysis (PCA) was generated using plotPCA. Volcano plots, heatmaps, and violin plots were created with ggplot2, EnhancedVolcano, pheatmap, and patchwork [51–54].

### Gene Set Enrichment Analysis

GSEA was performed in R using clusterProfiler. Genes were ranked by log₂ fold change from DESeq2 results and converted to Entrez IDs via org.Mm.eg.db. Enrichment was assessed for GO Biological Processes, KEGG, and Reactome pathways using gseGO, gseKEGG, and gsePathway. Pathways related to cardiac stress, DNA repair, genomic stability, autophagy, mitophagy, and immune processes were prioritized. Normalized enrichment scores (NES) and adjusted p-values were visualized as heatmaps using pheatmap, with significance indicated by stars (*p < 0.05, **p < 0.01, ***p < 0.001). Protein–protein interaction network and functional enrichment analysis was performed with STRING database [49].

### Metabolomics Analysis

Following in vivo normoxic and hypoxic exposure, heart tissues were snap-frozen and cryo-pulverized (n=3 animals per species per condition). Cryo-pulverized heart tissues were weighed, minced, and homogenized with steel beads in a methanol–acetonitrile–water (2:1:1) extraction solution containing internal standards (d-Valine and d-Phenylalanine), followed by protein precipitation at −20 °C and centrifugation to collect the metabolite-rich supernatant, which was dried under nitrogen and reconstituted in methanol–water before LC-MS analysis. Samples were analyzed on a Thermo Scientific Vanquish UHPLC system coupled to an Orbitrap Exploris 240 mass spectrometer using C18 chromatography, operating in both positive and negative ion modes with high-resolution MS1 and data-dependent MS2 acquisition. Untargeted metabolomics data processing was performed using Compound Discoverer 3.3, applying filtering, compound grouping, spectral matching (mzCloud, ChemSpider, HMDB), composition prediction, and QC normalization to identify and compare metabolites across experimental groups.

### Metabolomics Data Analysis

Metabolite data were analyzed using MetaboAnalyst [55]. Data were log-transformed and auto-scaled for normalization. Univariate analysis was performed using t-tests with FDR correction, and volcano plots and fold-change analysis identified significant metabolites.

### High-Resolution Respirometry and Mitochondrial H₂O₂ Production

Mitochondria were freshly isolated from heart tissues immediately following *in vivo* normoxic and hypoxic exposure in Blind mole-rats (BMRs) and mouse (*n* = 3 animals per species per condition). Dissected heart tissues of BMR and mouse were rinsed in cold PBS-EDTA, minced, and incubated with 1 mL trypsin on ice (30 min). Cells were pelleted (200g, 5 min), resuspended in IBM1 buffer (67 mM sucrose, 50 mM KCl, 50 mM Tris-HCl, 10 mM EDTA, 0.2% fatty-acid free BSA, pH 7.4), and centrifuged sequentially (700 g, 10 min; 8000 g, 10 min) to isolate mitochondria. The pellet was washed in IBM2 buffer (250 mM sucrose, 0.3 mM EGTA-Tris, 10 mM Tris-HCl, pH 7.4), and mitochondrial protein quantified via DC assay. Mitochondria isolated from normoxic and hypoxic mouse and BMR hearts were used to measure oxygen consumption and H_2_O_2_ production rate via the Oroboros-O2k FluoRespirometer instrument. Oxygen consumption and H_2_O_2_ production was measured with AmplexUltraRed fluorescence sensor with green filters that were inserted into the O2k chambers. According to the manufacturer’s instructions, air and H_2_O_2_ calibrations were conducted by adding the Mitochondria Respiration Buffer (120 mM sucrose, 50 mM KCl, 20 mM Tris–HCl, 4 mM KH_2_PO_4_, 2 mM MgCl_2_, 1 mM EGTA, 1 mg/ml fatty-acid-free BSA, pH 7.2) to the chambers. Isolated mitochondria were added depending on the amount determined with DC assay. SUIT-006-D048 protocol was followed for H_2_O_2_ production during forward electron transport. 5 mM Pyruvate, 2 mM Malate, 2.5 mM ADP, 5 nM Oligomycin, 0.5 CCCP (repeated for some samples) and 2.5 Antimycin A, along with 0.1 M H_2_O_2_ titration (thrice) were added to chambers. For Hydrogen Peroxide measurement during reverse electron transport, SUIT026-D064 protocol was followed, including the addition of 10 mM Succinate, 0.5 Rotenone, 2.5 mM ADP, 2.5 Antimycin A with thrice 0.1 H_2_O_2_ titration. Data analysis was conducted according to the excel templates offered by the manufacturer.

### Immunoblotting

Following *in vivo* normoxic and hypoxic exposure, snap-frozen heart tissues were processed for immunoblot analysis. Heart samples were diced and homogenized (1:10, w/v) in an ice-cold RIPA buffer supplemented with protease and phosphatase inhibitors. Lysates were incubated on ice for 30 min and clarified by centrifugation (13,000 × g, 10 min, 4 °C). Protein concentrations were determined using a BCA assay (Cat#23227, Thermo Scientific).

Equal amounts of protein (∼30 – 50 µg per lane) were separated by SDS–PAGE using 7–12% polyacrylamide or gradient gels and transferred onto nitrocellulose or PVDF membranes. Following transfer, membranes were washed three times for 5 min with TBST and blocked for 1 h at room temperature in 5% non-fat dry milk or bovine serum albumin (BSA) prepared in Tris-buffered saline containing 0.05% Tween-20 (TBST). Following blocking, membranes were incubated overnight at 4 °C with primary antibodies of p-AMPK, AMPK, p-mTOR, mTOR, p-ULK, LC3, Beclin-1, FUNDC1, caspase-3, GAPDH, and β-actin diluted at 1:500–1:1000 (see Table S1 for antibody details). Membranes were then washed three times with TBST and incubated for 1 h at room temperature with HRP-conjugated secondary antibodies (anti-rabbit or anti-mouse) diluted at 1:2000–1:5000 in 1% BSA or 1% non-fat dry milk/TBST solution. After final washes, immunoreactive bands were visualized using enhanced chemiluminescence and imaged with a ChemiDoc MP Imaging System (Bio-Rad). Band intensities were quantified using ImageJ software (NIH) and normalized to appropriate loading controls.

### Cardiomyocyte Isolation

Primary cardiomyocytes were isolated from adult male blind mole-rats (BMRs) and rats for use in in vitro experiments using a modified Langendorff perfusion protocol, as described previously (PMID: 21062918). Briefly, animals were anesthetized with isoflurane inhalation following intraperitoneal administration of heparin, with the dose adjusted according to body weight. All surgical instruments and the animals were disinfected with 70% ethanol. Hearts were rapidly excised with the aorta and immediately placed in a freshly prepared Ca²⁺-free solution. The aorta was cannulated and ligated with surgical suture, and hearts were perfused at 37 °C in a constant-pressure perfusion system with Ca²⁺-free buffer containing (in mmol/L): 137 NaCl, 5.7 KCl, 4.4 NaHCO₃, 1.5 KH₂PO₄, 3.6 MgCl₂, 20 HEPES, 10 glucose, and 20 taurine (pH 7.2), continuously bubbled with 100% oxygen. Hearts were initially perfused for 10 min to remove residual blood and debris.

Enzymatic digestion was performed by recirculating solution supplemented with 0.5 mg/mL collagenase type II (Cat#77336S, Cell Signaling) for 10–15 min at 37 °C with a constant speed. Following digestion, hearts were removed from the perfusion apparatus, transferred to collagenase-free solution containing 2 mg/mL bovine serum albumin (BSA), and mechanically dissociated into small pieces with scissors. Cardiomyocytes were extracted from the small tissues using a pipette and filtered through a 100-µm cell strainer. Cells were washed once with collagenase-free solution and allowed to sediment by gravity for 10 minutes. Following, calcium tolerance was restored by sequential washes in buffers containing increasing Ca²⁺ concentrations (0.3 mM, 0.5 mM, and 1.8 mM). This procedure yielded approximately 90% viable, rod-shaped cardiomyocytes. Isolated cardiomyocytes were in the HEPES-buffered solution containing (in mmol/L): 137 NaCl, 4 KCl, 1 MgCl₂, 10 HEPES, 1.8 CaCl₂, and 10 glucose (pH 7.4) at 37 °C standard incubator until used in downstream experiments.

### Hypoxia/Reoxygenation Treatment and Drug Administration

Primary cardiomyocytes and H9c2 cells (WT and ULK1 KI) were treated with Mdivi-1 (20 µM) or urolithin A (5 µM); primary cardiomyocytes were treated in Tyrode solution (137 mM NaCl, 4 mM KCl, 1 mM MgCl₂, 10 mM HEPES, 1.8 mM CaCl₂, 10 mM glucose, pH 7.4), and H9c2 cells were treated in DMEM (10% FBS, without penicillin/streptomycin). Vehicle controls received an equivalent volume of DMSO.

Primary cardiomyocytes were pretreated for 16 h, then subjected to hypoxia (1% O₂, 37°C) for 30 min followed by reoxygenation under normoxia (21% O₂, 37°C) for 2 h. H9c2 cells were pretreated for 24 h, then exposed to hypoxia (1% O₂, 37°C) for 24 h followed by reoxygenation under normoxia (21% O₂, 37°C) for 24 h. Hypoxia was performed in a hypoxia incubator; reoxygenation was performed in a standard incubator.

### Cell viability Assessment

Primary cardiomyocytes subjected to hypoxia–reoxygenation (H/R) with or without pharmacological treatment were evaluated by morphology-based counting. The identification of live and dead cardiomyocyte cells was based on the rod-shaped phenotype characteristic of cardiomyocytes. Brightfield images of the entire well were acquired before and after H/R. Cells were manually counted using ImageJ software. Cell viability was calculated as the percentage of viable cells relative to the total number of cells according to the following formula:

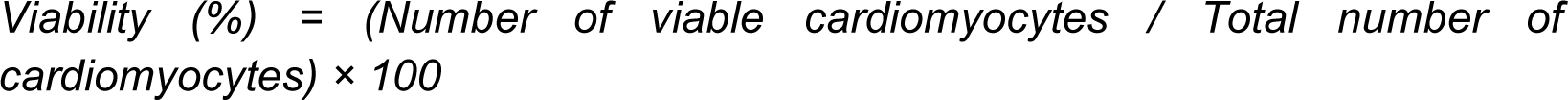

Cell viability in WT and ULK1 knock-in H9c2 cells was assessed using the MTS assay (CellTiter 96® AQueous One Solution, Promega) according to the manufacturer’s instructions. Absorbance was measured at 490 nm using a microplate reader. Data represent three independent experiments performed in technical triplicate, normalized to untreated controls within each genotype and expressed as mean ± SEM.

### Mitochondrial Morphology

Mitochondrial morphology was evaluated using MitoTracker Green FM (Cell Signaling Technology, Cat# 9074S). Briefly, MitoTracker Green FM was dissolved in DMSO to generate a 1 mM stock solution. Primary cardiomyocytes were seeded at equal densities in 96-well plates, treated as described above, and subjected to hypoxia/reoxygenation (H/R). Following H/R, cells were stained with MitoTracker Green FM according to the manufacturer’s instructions, and fluorescence images were acquired using a fluorescence microscope under identical exposure settings for all groups.

### Quantitative Analysis of Mitochondrial Morphology

Mitochondrial morphology was quantified using Fiji/ImageJ (v1.53). Fluorescence images were first processed using a median filter (radius = 1 pixel) to reduce salt-and-pepper noise, followed by background subtraction using the rolling ball algorithm (radius = 50 pixels). Images were converted to 8-bit format and segmented using Otsu automatic thresholding to generate binary masks. Particles smaller than 0.1 μm² were excluded to remove background noise and nonspecific signals.Individual mitochondrial structures were analyzed using the *Analyze Particles* plugin. The following parameters were quantified: mitochondrial area (μm²), aspect ratio (AR; major/minor axis), circularity (4π × area/perimeter²), and form factor (1/circularity), which reflects mitochondrial network complexity and branching. For each condition, approximately 48 cells were analyzed per biological replicate, yielding more than 1,400 individual mitochondrial structures per group. Images were acquired using either 40× (27.31 pixels/μm) or 10× (6.83 pixels/μm) objectives, and pixel-to-micron calibration was applied accordingly. Statistical analysis was performed as described in the figure legends [69].

### Multiple Sequence Alignment of ULK1 Orthologs

Multiple sequence alignments of ULK1 orthologs from human (O75385), mouse (O70405), rat (D3ZMG0), naked mole-rat (XP_012927320.1), and blind mole-rat (XP_029412313.1) were performed using Clustal Omega [71]. The resulting alignment was visualized and annotated to highlight species-specific variations in the kinase domain.

### Structural Modeling of Blind mole-rat ULK1

The kinase domain of BMR ULK1 was modeled using AlphaFold2 (AF2) [72] and AlphaFold3 (AF3) [73]. For each method, five predictions were performed and the top-ranking model based pLDDT was selected. For AF2, predictions were generated via ColabFold with three recycling rounds, and the final model was relaxed [74–77]. The human ULK1 kinase domain crystal structure (PDB ID: 8p5l), co-crystallized with a kinase inhibitor, was used as the structural reference. Predicted BMR models were aligned to the human structure to identify the BMR-specific insertion and evaluate its structural context.

Solvent accessible surface area (SASA) calculations were performed at both the overall and per-residue levels using the DSSP module in Biopython [78,79]. To ensure consistent residue coverage across all models, the predicted structures were trimmed to match the human ULK1 domain spanning residues 8–280. The missing internal segment (residues 149–154) in the crystal structure was reconstructed using the MODELLER plugin within ChimeraX [80]. All structural visualizations were rendered in ChimeraX [81]. The SASA profiles were plotted to assess differences in surface exposure, particularly in the region encompassing the BMR-specific insertion (residues 164–171). Root mean square displacements (RMSD) were calculated referencing the Cα trace of the PDB structure via ChimeraX.

### Homology-Directed Repair Design

To identify conserved target regions within the ULK1(Serine/threonine-protein kinase) gene, genomic alignments were performed across *Nannospalax galili* (Scaffold 566, Ref: NW_008355811.1), *Rattus norvegicus*, and *Homo sapiens*. Based on evolutionary conservation, an 8-amino acid insertion site was selected. Three target-specific crRNAs (ULK1_crRNA1–3) and a negative control sgRNA were designed using the Alt-R CRISPR-Cas9 Design Tool (Integrated DNA Technologies, IDT) to minimize off-target activity. The specific sgRNA sequences targeting the locus are given in Table S3. For homology-directed repair (HDR), double-stranded donor templates were synthesized (IDT) containing the desired 24-nucleotide insertion flanked by homology arms. Synonymous mutations were engineered into the donor sequence at the Protospacer Adjacent Motif (PAM) site to prevent re-cleavage by Cas9 post-integration.

### H9c2 Cell Culture

H9c2 rat cardiomyocyte cells (ATCC, CRL-1446) were maintained in Dulbecco’s Modified Eagle Medium (DMEM, Gibco) supplemented with 10% fetal bovine serum (FBS, Sigma-Aldrich) and 0.5% penicillin-streptomycin (Gibco) at 37°C in 5% CO₂. For maintenance (passages 9 to 21), cells were passaged every 4 days using 0.25% trypsin-EDTA (Gibco). Twenty-four hours prior to nucleofection, cells were seeded at a density of 1×10^5^ cells/mL.

### CRISPR–Cas9 Genome Editing and Knock-In Generation

Genome editing was performed via ribonucleoprotein (RNP) delivery. Cas9 (Alt-R EnGen S. pyogenes Cas9 NLS, NEB) was complexed with synthetic sgRNAs (IDT) for 10 min at room temperature. H9c2 cells were resuspended in nucleofection solution (1 × 10⁵ cells/mL), mixed with RNP complexes and 1 μg donor template, and electroporated using a 4D-Nucleofector system (Lonza; program DS-120). Cells were recovered in RPMI 1640 for 10 min and transferred to antibiotic-free DMEM containing 20% FBS. Editing efficiency was assessed 72 h post-transfection. Cells were passaged 1–2 times prior to clonal isolation.Monoclonal lines were generated by limiting dilution in 96-well plates. Monoclonality was verified by light microscopy. Clones were expanded in DMEM supplemented with 20% FBS and 0.25% penicillin–streptomycin for 7–10 days prior to genotyping.

### Genotyping and PCR

Genomic DNA was isolated from expanded clones using the Monarch® Genomic DNA Purification Kit (NEB #T3010) and quantified via NanoDrop spectrophotometry (Thermo Fisher Scientific). Target integration was verified by PCR using Q5® High-Fidelity DNA Polymerase (NEB #M0491). Reactions (25 μL) contained 50–100 ng genomic DNA, 200 μM dNTPs, 0.5 μM primers, and 0.5 U Q5 polymerase. Each PCR mixture was initially denatured at 98°C for 30 s and then cycled as follows: for Junction PCR, 30 cycles at 98°C for 10 s, 68–70°C for 20 s, and 72°C for 30 s; for Gradient PCR, 10 cycles at 98°C for 10 s, 70°C for 20 s, and 72°C for 12 s followed by 20 cycles at 98°C for 10 s, 64°C for 20 s, and 72°C for 12 s; with a final extension at 72°C for 2 min. PCR products were resolved on a 2% agarose gel with EZView Stain (A4205, BIOMATIC). Positive clones were validated by Sanger sequencing and BLAST analysis. The primer sets used in screening and sequencing are given in Table S3.

### ROS Measurement (H₂DCFDA Assay)

Intracellular reactive oxygen species (ROS) production was assessed using H₂DCFDA (2′,7′-dichlorodihydrofluorescein diacetate) in WT and ULK1 KI cardiomyocytes. Cells were cultured under standard conditions and grown to obtain 3 × 10⁶–4 × 10⁶ cells on the day prior to the experiment. Cells were subjected to normoxia (21% O₂) or hypoxia/reoxygenation (H/R; 24 h at 1% O₂, 5% CO₂, balance N₂, followed by 24 h reoxygenation at normoxia). After treatment, cells were plated in dark, clear-bottom 96-well plates (25,000 cells/well; n = 4 wells per condition) and allowed to adhere overnight.

For assay controls, H₂O₂ was prepared by diluting a 10 mM stock to 150 µM in 10% FBS DMEM/F12 and applied to cells for 30 min at 37 °C as a positive control; wells containing medium only were included as background controls. Cells were then incubated with H₂DCFDA (50 µM; 100 µL/well) for 45 min at 37 °C in the dark. The dye solution was removed and replaced with phenol red–free complete medium (10% FBS DMEM/F12), and fluorescence was measured immediately using a plate reader (Ex/Em 485/535 nm; endpoint mode).

### Cell Viability Assessment (CCK-8)

Following ROS fluorescence acquisition, cell viability was quantified in the same plate using the CCK-8 assay. Without removing the existing medium, CCK-8 reagent was added at a 1:5 dilution in 10% FBS DMEM/F12 (100 µL/well) and incubated for 2 h at 37 °C, 5% CO₂. Absorbance was measured at 450 nm using a microplate reader. ROS fluorescence values were normalized to the corresponding CCK-8 signal to account for differences in cell number/metabolic activity. Normalized values were expressed relative to the WT normoxia condition, which was set to 1.

### Fluorescence Imaging of Intracellular ROS

WT and ULK1 KI cells were subjected to normoxia or hypoxia/reoxygenation as described above. Cells were incubated with 50 μM H₂DCFDA for 45 min at 37 °C in the dark, washed with PBS to remove excess dye, and replenished with complete culture medium prior to live-cell imaging. Fluorescence images were acquired using an EVOS fluorescence microscope (10× objective). Fluorescence intensity was quantified using Fiji/ImageJ. Statistical analysis was performed using one-way ANOVA.

## Resource Availability

All data and code will be publicly available as of the date of publication.

## Acknowledgments

We thank all members of the Bozaykut-Eker lab and Esma Nur Okatan for helpful discussion. We also thank Zsuzsanna Ortutay for technical assistance with CRISPR optimization and Sanger sequencing analysis. This work was supported by TUBITAK-NSF Bilateral Program grant number 121N467. Metabolomics was performed by Szaomics Company (Istanbul, Turkey), transcriptomics was performed by Tufts Genomics Core (Boston, USA). P.B. received support from USDA cooperative agreement USDA/ARS 8050-10700-003.

## Author Contributions

All authors helped with designing the experiments. D.Y., S.F.O., E.O.A., C.K.O., H.F., I.S.Y, P.B., E.T., S.A.D., E.T. collected data and contributed to data analysis. F.C. performed the field study. D.Y. and P.B. analyzed data and wrote the paper with contributions from all other authors. All authors reviewed and approved the final manuscript.

## References

[1] Lee P, Chandel NS, Simon MC. Cellular adaptation to hypoxia through hypoxia-inducible factors and beyond. Nat Rev Mol Cell Biol. 2020;21(5):268–283. doi:10.1038/s41580-020-0216-1.

[2] Li H, Xiao F, Zhou C, Zhu T, Wang S. Metabolic adaptations and therapies in cardiac hypoxia: mechanisms and clinical implications/potential strategies. JACC Basic Transl Sci. 2025;10(6):862–878. doi:10.1016/j.jacbts.2024.11.009.

[3] Coimbra-Costa D, Alva N, Duran M, Carbonell T, Rama R. Oxidative stress and apoptosis after acute respiratory hypoxia and reoxygenation in rat brain. Redox Biol. 2017;12:216– 225. doi:10.1016/j.redox.2017.02.024.

[4] Wanderer AA. Hypoxia and inflammation. N Engl J Med. 2011;364(20):1976–1977. doi:10.1056/NEJMc1102876.

[5] Li F, Qiao Z, Duan Q, Nevo E. Adaptation of mammals to hypoxia. Anim Models Exp Med. 2021;4(4):311–318. doi:10.1002/ame2.12191.

[6] Eaton L, Pamenter ME. What to do with low O2: redox adaptations in vertebrates native to hypoxic environments. Comp Biochem Physiol A Mol Integr Physiol. 2022;271:111259. doi:10.1016/j.cbpa.2022.111259.

[7] Schmidt H, Malik A, Bicker A, Poetzsch G, Avivi A, Shams I, et al. Hypoxia tolerance, longevity and cancer-resistance in the mole rat Spalax: a liver transcriptomics approach. Sci Rep. 2017;7(1):14348. doi:10.1038/s41598-017-14013-2.

[8] Lagunas-Rangel FA. Cancer-free aging: insights from Spalax ehrenbergi superspecies. Ageing Res Rev. 2018;47:18–23. doi:10.1016/j.arr.2018.06.001.

[9] Malik A, Domankevich V, Han L, Fang X, Korol A, Avivi A, et al. Genome maintenance and bioenergetics of the long-lived hypoxia-tolerant and cancer-resistant blind mole-rat, Spalax: a cross-species analysis of brain transcriptome. Sci Rep. 2016;6:38624. doi:10.1038/srep38624.

[10] Widmer HR, Hoppeler H, Nevo E, Taylor CR, Weibel ER. Working underground: respiratory adaptations in the blind mole-rat. Proc Natl Acad Sci U S A. 1997;94(5):2062–2067. doi:10.1073/pnas.94.5.2062.

[11] Inci N, Akyildiz EO, Bulbul AA, Turanli ET, Akgun E, Baykal AT, et al. Transcriptomics and proteomics analyses reveal JAK signaling and inflammatory phenotypes during cellular senescence in blind mole-rats: the reflections of superior biology. Biology (Basel). 2022;11(9):1293. doi:10.3390/biology11091293.

[12] Inci N, Kamali D, Akyildiz EO, Tahir Turanli E, Bozaykut P. Translation of cellular senescence to novel therapeutics: insights from alternative tools and models. Front Aging. 2022;3:955275. doi:10.3389/fragi.2022.955275.

[13] Jang JY, Blum A, Liu J, Finkel T. The role of mitochondria in aging. J Clin Invest. 2018;128(9):3662–3670. doi:10.1172/JCI120842.

[14] Okoye CN, Koren SA, Wojtovich AP. Mitochondrial complex I ROS production and redox signaling in hypoxia. Redox Biol. 2023;67:102715. doi:10.1016/j.redox.2023.102715.

[15] Marchi S, Giorgi C, Suski JM, Agnoletto C, Bononi A, Bonora M, et al. Mitochondria-ROS crosstalk in the control of cell death and aging. J Signal Transduct. 2012;2012:329635. doi:10.1155/2012/329635.

[16] Xu X, Pang Y, Fan X. Mitochondria in oxidative stress, inflammation and aging: from mechanisms to therapeutic advances. Signal Transduct Target Ther. 2025;10(1):190. doi:10.1038/s41392-025-02253-4.

[17] Chen S, Li Q, Shi H, Li F, Duan Y, Guo Q. New insights into the role of mitochondrial dynamics in oxidative stress-induced diseases. Biomed Pharmacother. 2024;178:117084. doi:10.1016/j.biopha.2024.117084.

[18] Gottlieb RA, Thomas A. Mitophagy and mitochondrial quality control mechanisms in the heart. Curr Pathobiol Rep. 2017;5(2):161–169. doi:10.1007/s40139-017-0133-y.

[19] Diao RY, Gustafsson ÅB. Mitochondrial quality surveillance: mitophagy in cardiovascular health and disease. Am J Physiol Cell Physiol. 2022;322(2):C218–C230. doi:10.1152/ajpcell.00360.2021.

[20] Titus AS, Sung EA, Zablocki D, Sadoshima J. Mitophagy for cardioprotection. Basic Res Cardiol. 2023;118(1):42. doi:10.1007/s00395-023-01009-x.

[21] Xu S, Wang Z, Guo F, Zhang Y, Peng H, Zhang H, et al. Mitophagy in ischemic heart disease: molecular mechanisms and clinical management. Cell Death Dis. 2024;15(1):1. doi:10.1038/s41419-024-06412-3.

[22] Burkewitz K, Zhang Y, Mair WB. AMPK at the nexus of energetics and aging. Cell Metab. 2014;20(1):10–25. doi:10.1016/j.cmet.2014.03.002.

[23] Wu CW, Storey KB. mTOR signaling in metabolic stress adaptation. Biomolecules. 2021;11(5):681. doi:10.3390/biom11050681.

[24] Weichhart T. mTOR as regulator of lifespan, aging, and cellular senescence: a mini-review. Gerontology. 2018;64(2):127–134. doi:10.1159/000477539.

[25] Park JM, Lee DH, Kim DH. Redefining the role of AMPK in autophagy and the energy stress response. Nat Commun. 2023;14(1):2994. doi:10.1038/s41467-023-38401-z.

[26] Zhang H, Liu B, Li T, Zhu Y, Luo G, Jiang Y, et al. AMPK activation serves a critical role in mitochondria quality control via modulating mitophagy in the heart under chronic hypoxia. Int J Mol Med. 2018;41(1):69–76. doi:10.3892/ijmm.2017.3213.

[27] Gui D, Cui Z, Zhang L, Yu C, Yao D, Xu M, et al. Salidroside attenuates hypoxia-induced pulmonary arterial smooth muscle cell proliferation and apoptosis resistance by upregulating autophagy through the AMPK–mTOR–ULK1 pathway. BMC Pulm Med. 2017;17(1):191. doi:10.1186/s12890-017-0477-4.

[28] Park TJ, Reznick J, Peterson BL, Blass G, Omerbašić D, Bennett NC, et al. Fructose-driven glycolysis supports anoxia resistance in the naked mole-rat. Science. 2017;356(6335):307–311. doi:10.1126/science.aab3896.

[29] Yim WWY, Yamamoto H, Kono K, et al. A pulse-chasable reporter processing assay for mammalian autophagy monitoring. eLife. 2022;11:e78923. doi:10.7554/eLife.78923.

[30] Tabata Fukushima C, Dancil IS, Clary H, Shah N, Nadtochiy SM, Brookes PS. Reactive oxygen species generation by reverse electron transfer at mitochondrial complex I under simulated early reperfusion conditions. Redox Biol. 2024;70:103047. doi:10.1016/j.redox.2024.103047.

[31] Liang H, Song K. Comprehensive metabolomics and transcriptomics analysis reveals protein and amino acid metabolic characteristics in liver tissue under chronic hypoxia. PLoS One. 2023;18(9):e0291698. doi:10.1371/journal.pone.0291698.

[32] Tang L, Zeng J, Geng P, Fang C, Wang Y, Sun M, et al. Global metabolic profiling identifies a pivotal role of proline and hydroxyproline metabolism in supporting hypoxic response in hepatocellular carcinoma. Clin Cancer Res. 2018;24(2):474–485. doi:10.1158/1078-0432.CCR-17-1707.

[33] Burke L, Guterman I, Palacios Gallego R, Britton RG, Burschowsky D, Tufarelli C, et al. The Janus-like role of proline metabolism in cancer. Cell Death Discov. 2020;6(1):104. doi:10.1038/s41420-020-00341-8.

[34] Phang JM. Proline metabolism in cell regulation and cancer biology: recent advances and hypotheses. Antioxid Redox Signal. 2019;30(4):635–649. doi:10.1089/ars.2017.7350.

[35] Pandhare J, Donald SP, Cooper SK, Phang JM. Regulation and function of proline oxidase under nutrient stress. J Cell Biochem. 2009;107(4):759–768. doi:10.1002/jcb.22174.

[36] Bai D, Zhou Y, Jing L, Guo C, Yang Q. Arginine metabolism in cancer biology and immunotherapy. Immune Netw. 2025;25(4):e30. doi:10.4110/in.2025.25.e30.

[37] Assi G, Faour WH. Arginine deprivation as a treatment approach targeting cancer cell metabolism and survival: a review of the literature. Eur J Pharmacol. 2023;953:175838. doi:10.1016/j.ejphar.2023.175838.

[38] Wang D, Duan JJ, Guo YF, Chen JJ, Chen TQ, Wang J, et al. Targeting the glutamine-arginine-proline metabolism axis in cancer. J Enzyme Inhib Med Chem. 2024;39(1):2367129. doi:10.1080/14756366.2024.2367129.

[39] Wang J, Xue Z, Hua C, Lin J, Shen Z, Song Y, et al. Metabolomic analysis of the ameliorative effect of enhanced proline metabolism on hypoxia-induced injury in cardiomyocytes. Oxid Med Cell Longev. 2020;2020:9141906. doi:10.1155/2020/9141906.

[40] Eisenberg T, Abdellatif M, Schroeder S, Primessnig U, Stekovic S, Pendl T, et al. Cardioprotection and lifespan extension by the natural polyamine spermidine. Nat Med. 2016;22(12):1428–1438. doi:10.1038/nm.4222.

[41] Zhang H, Wang J, Li L, Chai N, Chen Y, Wu F, et al. Spermine and spermidine reversed age-related cardiac deterioration in rats. Oncotarget. 2017;8(39):64793–64808. doi:10.18632/oncotarget.18334.

[42] Chai N, Zhang H, Li L, Yu X, Liu Y, Lin Y, et al. Spermidine prevents heart injury in neonatal rats exposed to intrauterine hypoxia by inhibiting oxidative stress and mitochondrial fragmentation. Oxid Med Cell Longev. 2019;2019:5406468. doi:10.1155/2019/5406468.

[43] Hofer SJ, Simon AK, Bergmann M, Eisenberg T, Kroemer G, Madeo F. Mechanisms of spermidine-induced autophagy and geroprotection. Nat Aging. 2022;2(12):1112–1129. doi:10.1038/s43587-022-00322-9.

[44] Wang J, Li S, Wang J, Wu F, Chen Y, Zhang H, et al. Spermidine alleviates cardiac aging by improving mitochondrial biogenesis and function. Aging (Albany NY). 2020;12(23):23693–23712. doi:10.18632/aging.102647.

[45] Qi Y, Qiu Q, Gu X, Tian Y, Zhang Y. ATM mediates spermidine-induced mitophagy via PINK1 and Parkin regulation in human fibroblasts. Sci Rep. 2016;6:24700. doi:10.1038/srep24700.

[46] Sözen M. A biological investigation on Turkish Spalax Guldentaedt, 1770 (Mammalia: Rodentia). G. U. J. Sci. 2005;18(2):167–181.

[47] Ewels PA, Peltzer A, Fillinger S, Patel H, Alneberg J, Wilm A, Garcia MU, Di Tommaso P, Nahnsen S. The nf-core framework for community-curated bioinformatics pipelines. Nat Biotechnol. 2020;38(3):276–278. doi:10.1038/s41587-020-0439-x.

[48] Love MI, Huber W, Anders S. Moderated estimation of fold change and dispersion for RNA-seq data with DESeq2. Genome Biol. 2014;15(12):550. doi:10.1186/s13059-014-0550-8.

[49] Szklarczyk D, Kirsch R, Koutrouli M, Nastou K, Mehryary F, Hachilif R, Gable AL, Fang T, Doncheva NT, Pyysalo S, Bork P, Jensen LJ, von Mering C. The STRING database in 2023: protein-protein association networks and functional enrichment analyses for any sequenced genome of interest. Nucleic Acids Res. 2023;51(D1):D638–D646. doi:10.1093/nar/gkac1000.

[50] Luo W, Brouwer C. Pathview: an R/Bioconductor package for pathway-based data integration and visualization. Bioinformatics. 2013;29(14):1830–1831. doi:10.1093/bioinformatics/btt285.

[51] Wickham H. ggplot2: Elegant graphics for data analysis. New York: Springer-Verlag; 2016.

[52] Blighe K, Rana S, Lewis M. EnhancedVolcano: publication-ready volcano plots with enhanced colouring and labeling. R package version 1.29.1. 2025. doi:10.18129/B9.bioc.EnhancedVolcano.

[53] Kolde R. pheatmap: Pretty heatmaps. R package version 1.0.13. 2025.

[54] Pedersen TL. patchwork: The composer of plots. R package version 1.3.2.9000. 2025.

[55] Pang Z, Lu Y, Zhou G, Hui F, Xu L, Viau C, Spigelman A, MacDonald P, Wishart D, Li S, Xia J. MetaboAnalyst 6.0: towards a unified platform for metabolomics data processing, analysis and interpretation. Nucleic Acids Res. 2024;52(W1):W398–W406. doi:10.1093/nar/gkae253.

[56] Di Tommaso P, Chatzou M, Floden EW, Barja PP, Palumbo E, Notredame C. Nextflow enables reproducible computational workflows. Nat Biotechnol. 2017;35(4):316–319. doi:10.1038/nbt.3820.

[57] Andrews S. FastQC: a quality control tool for high throughput sequence data. 2010. Available from: https://www.bioinformatics.babraham.ac.uk/projects/fastqc/.

[58] Martin M. Cutadapt removes adapter sequences from high-throughput sequencing reads. EMBnet.journal. 2011;17(1):10–12. doi:10.14806/ej.17.1.200.

[59] Patro R, Duggal G, Love MI, Irizarry RA, Kingsford C. Salmon provides fast and bias-aware quantification of transcript expression. Nat Methods. 2017;14(4):417–419. doi:10.1038/nmeth.4197.

[60] Love MI, Soneson C, Patro R. Swimming downstream: statistical analysis of differential transcript usage following Salmon quantification. F1000Res. 2018;7:952. doi:10.12688/f1000research.15398.3.

[61] Okonechnikov K, Conesa A, García-Alcalde F. Qualimap 2: advanced multi-sample quality control for high-throughput sequencing data. Bioinformatics. 2016;32(2):292–294. doi:10.1093/bioinformatics/btv566.

[62] Wang L, Wang S, Li W. RSeQC: quality control of RNA-seq experiments. Bioinformatics. 2012;28(16):2184–2185. doi:10.1093/bioinformatics/bts356.

[63] Sayols S, Scherzinger D, Klein H. dupRadar: a Bioconductor package for the assessment of PCR artifacts in RNA-Seq data. BMC Bioinformatics. 2016;17(1):428. doi:10.1186/s12859-016-1276-2.

[64] Dobin A, Davis CA, Schlesinger F, Drenkow J, Zaleski C, Jha S, Batut P, Chaisson M, Gingeras TR. STAR: ultrafast universal RNA-seq aligner. Bioinformatics. 2013;29(1):15– 21. doi:10.1093/bioinformatics/bts635.

[65] Liao Y, Smyth GK, Shi W. featureCounts: an efficient general purpose program for assigning sequence reads to genomic features. Bioinformatics. 2014;30(7):923–930. doi:10.1093/bioinformatics/btt656.

[66] Broad Institute. Picard toolkit. Broad Institute, GitHub repository. 2019. Available from: https://broadinstitute.github.io/picard/.

[67] Pertea M, Pertea GM, Antonescu CM, Chang TC, Mendell JT, Salzberg SL. StringTie enables improved reconstruction of a transcriptome from RNA-seq reads. Nat Biotechnol. 2015;33(3):290–295. doi:10.1038/nbt.3122.

[68] Li H, Handsaker B, Wysoker A, Fennell T, Ruan J, Homer N, Marth G, Abecasis G, Durbin R; 1000 Genome Project Data Processing Subgroup. The Sequence Alignment/Map format and SAMtools. Bioinformatics. 2009;25(16):2078–2079. doi:10.1093/bioinformatics/btp352.

[69] Quinlan AR, Hall IM. BEDTools: a flexible suite of utilities for comparing genomic features. Bioinformatics. 2010;26(6):841–842. doi:10.1093/bioinformatics/btq033.

[70] Valente AJF, Maddalena LA, Robb EL, Moradi F, Stuart JA. A simple ImageJ macro tool for analyzing mitochondrial network morphology in mammalian cell culture. Acta Histochem. 2017;119(3):315–326. doi:10.1016/j.acthis.2017.03.001.

[71] Sievers F, Higgins DG. Clustal Omega for making accurate alignments of many protein sequences. Protein Sci. 2018;27(1):135–145. doi:10.1002/pro.3290.

[72] Jumper J, Evans R, Pritzel A, Green T, Figurnov M, Ronneberger O, Tunyasuvunakool K, et al. Highly accurate protein structure prediction with AlphaFold. Nature. 2021;596(7873):583–589. doi:10.1038/s41586-021-03819-2.

[73] Abramson J, Adler J, Dunger J, Evans R, Green T, Pritzel A, Ronneberger O, et al. Accurate structure prediction of biomolecular interactions with AlphaFold 3. Nature. 2024;630(8017):493–500. doi:10.1038/s41586-024-07487-w.

[74] Mirdita M, Schütze K, Moriwaki Y, Heo L, Ovchinnikov S, Steinegger M. ColabFold: making protein folding accessible to all. Nat Methods. 2022;19(6):679–682. doi:10.1038/s41592-022-01488-1.

[75] Mirdita M, Steinegger M, Söding J. MMseqs2 desktop and local web server app for fast, interactive sequence searches. Bioinformatics. 2019;35(16):2856–2858. doi:10.1093/bioinformatics/bty1057.

[76] Mirdita M, von den Driesch L, Galiez C, Martin MJ, Söding J, Steinegger M. Uniclust databases of clustered and deeply annotated protein sequences and alignments. Nucleic Acids Res. 2017;45(D1):D170–D176. doi:10.1093/nar/gkw1081.

[77] Eastman P, Swails J, Chodera JD, McGibbon RT, Zhao Y, Beauchamp KA, Wang LP, Simmonett AC, Harrigan MP, Stern CD, et al. OpenMM 7: rapid development of high performance algorithms for molecular dynamics. PLoS Comput Biol. 2017;13(7):e1005659. doi:10.1371/journal.pcbi.1005659.

[78] Zacharias J, Knapp EW. Protein secondary structure classification revisited: processing DSSP information with PSSC. J Chem Inf Model. 2014;54(7):2166–2179. doi:10.1021/ci5000856.

[79] Cock PJA, Antao T, Chang JT, Chapman BA, Cox CJ, Dalke A, Friedberg I, Hamelryck T, Kauff F, Wilczynski B, de Hoon MJL. Biopython: freely available Python tools for computational molecular biology and bioinformatics. Bioinformatics. 2009;25(11):1422–1423. doi:10.1093/bioinformatics/btp163.

[80] Webb B, Sali A. Protein structure modeling with MODELLER. Methods Mol Biol. 2021;2199:239–255. doi:10.1007/978-1-0716-0892-0_14.

[81] Pettersen EF, Goddard TD, Huang CC, Couch GS, Greenblatt DM, Meng EC, Ferrin TE. UCSF ChimeraX: structure visualization for researchers, educators, and developers. Protein Sci. 2021;30(1):70–82. doi:10.1002/pro.3943.

